# Phosphorylation regulated conformational diversity and topological dynamics of an intrinsically disordered nuclear receptor

**DOI:** 10.1101/2024.09.21.614239

**Authors:** Vasily Akulov, Alba Jiménez Panizo, Eva Estebanez Perpiña, John van Noort, Alireza Mashaghi

**Affiliations:** Medical Systems Biophysics and Bioengineering, Division of Systems Pharmacology and Pharmacy, Leiden Academic Centre for Drug Research, Leiden University, 2333 CC, Leiden, the Netherlands; Leiden Institute of Physics, Leiden University, 2300 RA, Leiden, the Netherlands; Centre for Interdisciplinary Genome Research, Faculty of Science, Leiden University, 2333 CC, Leiden, the Netherlands; Department of Biochemistry and Molecular Biomedicine, Institute of Biomedicine (IBUB) of the University of Barcelona (UB), 08014, Barcelona, Spain

**Keywords:** glucocorticoid receptor, nuclear receptor, phosphorylation, post-translational modification, circuit topology, molecular dynamics simulations, intrinsically disordered protein

## Abstract

Site-specific phosphorylation of disordered proteins is often considered as a marker of protein activity, yet it is unclear how phosphorylation alters conformational dynamics of disordered protein chains, such as those in the nuclear receptor superfamily. In the case of disordered human glucocorticoid receptor N-terminal domain (GR NTD), a negatively charged region known as core activation function 1 (AF1c) features three phosphorylation sites, regulating its function and intracellular localization. Deletion of this sequence reduces GR transcriptional activation ability dramatically in cell experiments. By developing a circuit topology-based fold analysis approach, combined with atomistic simulations, we reveal that site-specific phosphorylation facilitates formation of non-local contacts, leading to the emergence of disordered compact topologies with significant entanglement, which are distinct from solvent exposed topologies. While we observe that the topological buildup of solvent-exposed states is similar in different phosphovariants, it depends on the exact phosphorylation site for the disordered compact states. This study thus reveals the complex regulatory role of the GR phosphorylation and introduces a unique analysis framework that can be broadly applied to studying topological dynamics of disordered proteins.

## Introduction

Protein molecules can adopt a broad spectrum of conformational states, ranging from disordered conformations to remarkably stable amyloid fibers ^1-3^. In contrast to stably folded proteins, which tend to form unique, dense globules stabilized by interchain contacts, disordered proteins exhibit dynamic, solute-exposed conformations. The absence of a dense globule leads to significant structural fluctuations in disordered proteins. However, these are extreme cases; many proteins lie somewhere in between. Folded proteins can contain dynamic, disordered loops and turns, and intrinsically disordered proteins (IDPs) can expectedly be found in compact, disordered states, where certain residues form transient contacts and thus the chain forms a highly connected yet dynamic plexus. Evidence suggests that “disordered compact” states can be observed in isolated IDPs depending on their sequence ^4^,are linked to function ^3,5^ and may be modulated by post-translational modifications. Reversible phosphorylation, for example, is thought to be a mechanism nature uses to modulate IDP function, presumably by regulating the conformational dynamics and compaction of these proteins. Consequently, IDPs presumably adopt a wide range of shapes and interact with different binding partners or partition to different subcellular spaces in a context-dependent manner ^6^, through largely unknown structural mechanisms. Despite active research on IDPs, these types of protein states have not been thoroughly studied, due to technical challenges.

Our understanding of IDPs is hindered by the lack of a proper description of the dynamics that capture topological motifs hidden within conformational noise. We do not yet know what patterns the transient contacts and dynamic loopy topologies form in disordered compact states, and whether these patterns can be used to identify these states. Recently, the topology of protein chains has been defined based on the arrangement of loops or the associated intrachain contacts. This approach, known as circuit topology (CT) ^7^, has been applied to stable folded proteins for various applications ^8,9^ and has proven effective for modeling polymer folding reactions ^10^. The approach has also been applied to disordered proteins, enabling the capture of conserved features in their topological fluctuations ^11^. Additionally, it has been used to detect topological similarities between IDPs with similar functions, providing a new metric for quantifying structural similarity suitable for both IDPs and proteins with stable 3D structures ^11^. As such, one may envision this method being applied to study phosphorylation-induced order in disordered chains, aiding in the identification and study of disordered compact conformations.

In this research, we develop an analysis framework by combining extensive molecular dynamics (MD) simulations and CT to study phosphorylation-regulated structural and dynamic properties of a disordered protein. We achieve this by studying the disordered N-terminal domain of the human glucocorticoid receptor (GR NTD) (Figure 1). This protein is a key transcription factor and a member of the nuclear receptor superfamily, which is implicated in many physiological and pathological processes such as metabolism, homeostasis and inflammation ^12^. The segment of the NTD, known as activation function 1 (AF1; comprising the residue region 77-262) and, more specifically, the AF1 core (AF1c, including residues 187-244) is of particular importance, as is the case in many other related nuclear receptors. The hormone-independent AF1 is required for maximal transcriptional activation of GR ^13^ . Deletion of this sequence reduces GR transcription activation ability for GR-dependent test genes by at least 60–70% ^14^. AF1 is subject to post-translational modifications, that contribute to the complexity and context-dependence of GR signaling ^15,16^. Site-specific phosphorylation of GR NTD is often considered a marker of protein activation ^14,17-26^ and a determinant of its subcellular localization. However, its implications for secondary or higher-order structural changes are largely unknown despite recent advances ^18,27-30^. Unfortunately, there is no detailed experimental structure neither for the full-length, nor for NTD fragments such as AF1c. Modeling data on its dynamics is also lacking, as the intrinsically disordered nature of AF1c precludes conformational analysis. Our IDP analysis approach reveals that phosphorylation tunes the structural diversity of this disordered protein and leads to the emergence of disordered compact states, which are detectable using the CT-based contact analysis approach.

**Figure 1:**
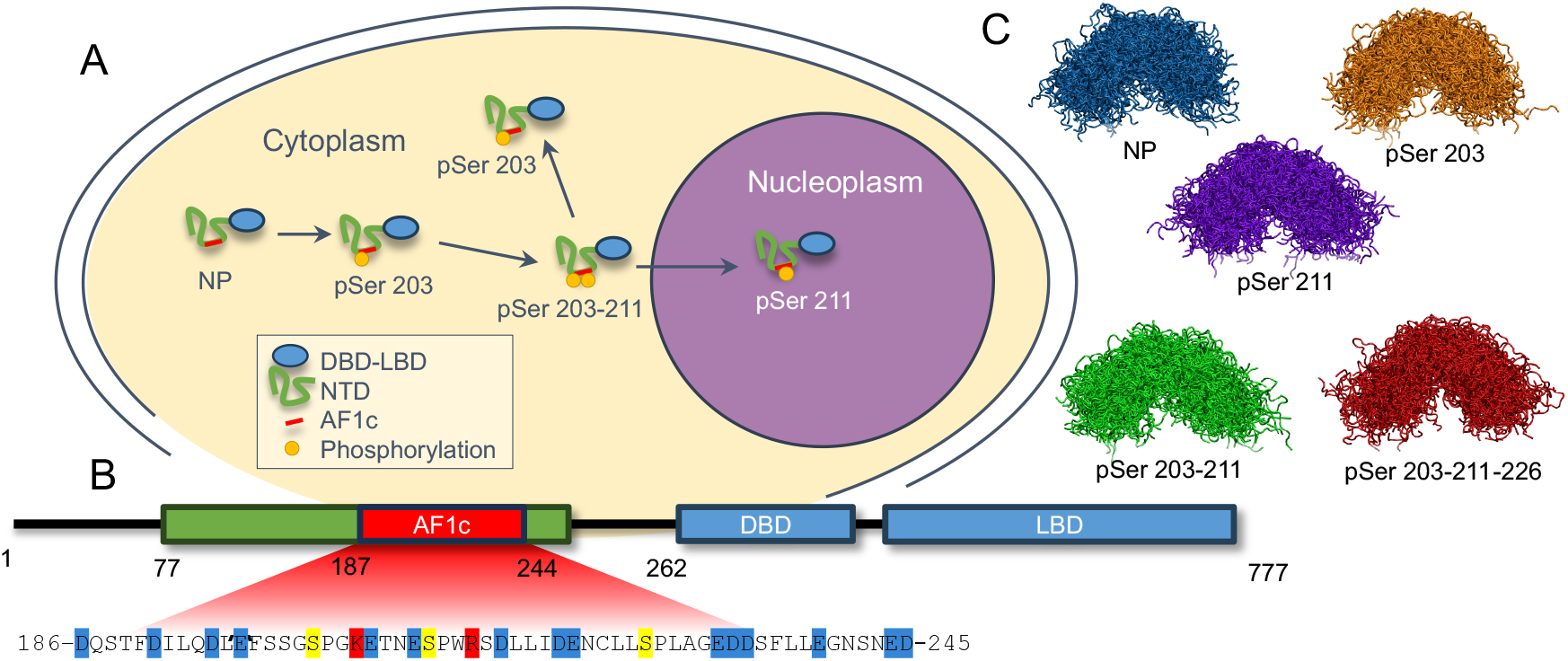
Role of phosphorylation in GR structure-function relationship. (**A**) The GR phosphorylation pathway. In non-activated form GR is not phosphorylated, but upon activation first get phosphorylated Ser 203, then Ser 211. After that, protein is transported to the nucleus, and loses Ser203 phosphorylation. Taken with changes from ^21^ (**B**) The schematic representation of GR with the sequence of AF1c region. GR has a modular architecture consisting of an intrinsically disordered amino-terminal domain (NTD; colored in green), including core activation function region 1 (AF1c in red), a zinc-finger DNA-binding domain (DBD), and a carboxy-terminal ligand-binding domain (LBD), each contributing to the receptor’s function. In the AF1c sequence, residues are color-coded: blue – negatively charged residues at neutral pH; red – positively charged residues at neutral pH; yellow – serine residues, which could be phosphorylated. (**C**) Overlaid structures of simulated AF1c phosphovariants.

## Methods

### Molecular dynamics simulations

For conventional MD simulations of AF1c, initial structures were predicted with LocalColabFold program ^31-33^. Because LocalColabFold does not recognize phosphoserine amino acids, it was replaced with glutamic acid in the input sequence of phosphorylated proteins. Then, after obtaining 3D-structures, these glutamic residues were replaced by the phosphoserines with the help of SwissSidechain v.2 PyMOL Plugin ^34-38^. The residue SEP2 was used. It has charge -2, that phosphoserine has at native pH conditions. The structures with the highest quality scores for each phosphorylation variant were used as initial structures. Three different initial structures were selected for simulating each of the phosphorylation variants.

The MD simulations were performed using the free molecular simulation software GROMACS^39^ version 2021.7 patched with the open-source, community-developed PLUMED library ^40^ version 2.8.3 ^41^. The protein force field was amber99SBdisp ^42,43^, with explicit solvent - amber99SBdisp water model, phosphoserine parameters were taken from ^44^. The AF1c was solvated in dodecahedron box with the box vector lengths 9.5 nm 9.5 nm 9.5 nm and periodic boundary conditions. The system was neutralized with 150 mM sodium and chloride ions. Before the production ran, the system was minimized and equilibrated. The equilibration was done in NVT and then NPT ensembles for 100 ps, the heavy protein atoms were restrained by harmonic force with the default value for the constants being 1000 kJ mol^-1^ nm^-2^. The LINCS algorithm ^45^ was used to constrain H-bonds vibrations. Integration was done with leap-frog algorithm, time step 2 fs. Buffered neighbor searching was done with Verlet list cutoff scheme, cutoff 1.0 nm in equilibration steps and 0.9 nm at the production step. Particle Mesh Ewald ^46^ with cubic interpolation was used for long-range electrostatics. In simulations, the pressure of 1 bar was controlled with a Parrinello-Rahman barostat, relaxation time 2 ps for equilibration steps and 5 ps for the production step. The isothermal compressibility of water was 4.5×10-5 bar^-1^. The solute and solvent were coupled independently; the temperature was set for both on 310 K. During equilibration steps, a modified Berendsen thermostat was used with a time constant tau of 0.1 ps. In production runs, the temperature was controlled by a Nose-Hoover thermostat, with time constant tau equal to 1 ps. The length of the simulation in production runs was 1000 ns for all systems. Eight replicates for each phosphorylation type were independently prepared and simulated. For analysis, the last 900 ns of simulations, with time steps of 500 ps were used. By removing the first 100 ns, we ensure that our simulation is not dependent on the choice of our initial conformation, as evidenced by the correlation time scales seen in our study. This part of the trajectory was considered as equilibrated. All frames for which the distance between the protein molecule and its periodic image was less than 2 nm were removed from the analysis. All trajectories for the same phosphorylation type were combined for the analysis.

### Data analysis

All analysis and plots preparation were done with homemade Python 3 scripts in the Anaconda based environments with help of different libraries: scipy ^47^, matplotlib ^48^, numpy ^49^, mdtraj ^50^, mdanalysis ^51-53^, pyemma ^54^, seaborn ^55^, pandas ^56^.

### Calculation of NMR spectra

The NMR shifts from simulated trajectories were calculated with SPARTA+ program ^57^. The experimental NMR spectrum was taken from ^58^. The NP sequence, used in this research, covers residues from 187 to 244 of GR, while in NMR research of *Kim et al*. the sequence ranges from 181 to 244 a.a. Correction of NMR backbone shift was done by substruction of the random coil component from both predicted and calculated spectra. This component was calculated with the help of Poulsen IDP/IUP random coil chemical shifts server ^59,60^. The secondary structure was computed with the help of the DSSP program ^61^. Protein structure images were generated with VMD package ^62^, (http://www.ks.uiuc.edu/Research/vmd/).

### Circuit Topology

For circuit topology analysis, trajectories were subsampled with a 2 ns time step to speed up the analysis. Next, these frames were saved as pdb files and transformed to CT representation with the code ProteinCT ^63^. This representation was then analyzed using newly developed code that can be found in GitHub repository: https://github.com/circuittopology/CT_Folding_Score.

## Results

### All-atom molecular dynamics simulations reveal the disordered nature of GR-AF1c under various phosphorylation states

We selected five phosphovariants of AF1c, which are reportedly visited by the GR protein as it transitions from the cytoplasm to the nucleus, to study the effect of individual phosphorylations on the ensemble of conformations (Figure 1A). The chain may get phosphorylated at three serine residues, namely residues 203, 211, and 226 (Figure 1B), which are followed by prolines in the primary sequence. The phosphorylation of these sites correlates with GR activation and localization as depicted in Figure 1A ^18-21,24,25^. The overlaid conformational samples of the simulated AF1c variants are shown in Figure 1C (see Methods). These phosphorylation variants are: non-phosphorylated variant (NP); AF1c with phosphorylated serine 203 (pSer 203); AF1c with phosphorylated serine 211 (pSer 211); AF1c with phosphorylated serine residues 203 and 211 (pSer 203-211); AF1c with phosphoserines at positions 203, 211 and 226 (pSer 203-211-226). All these phosphorylation variants, except pSer 203-211-226 were directly observed *in vivo* ^23^. The triple phosphorylated variant was observed *in vitro* ^25^. Here, we focus our research on AF1c with all prolines in the trans isomer, which is commonly seen in cells and leave the analysis of trans-to-cis isomerization to future studies.

During the course of simulations, we observed that all variants were showing significant conformational disorder (Figure 1C). As can be seen in Figure 1C, overlaid structures of different phosphovariants representing the distribution of monomers are characteristic of a disordered chain. Additionally, the high fluctuations in solvent accessible surface area (SASA) (Figure S2) per residue show the dynamic nature of the proteins studied. The disordered nature of AF1c can be attributed to the unfavorable folding balance between charges (−13 in non-phosphorylated case at neutral pH) and hydrophobicity.

### NMR shifts of simulated protein correlate well to experimentally measured values

From MD data of the NP variant, we calculated NMR chemical shifts using SPARTA+ ^57^ and compared them with available experimental NMR chemical shifts ^58^. Eight independent trajectories of the NP variant (8 microseconds in total) were combined for this analysis and the predicted NMR shift was correlated to the experimental data using the Pearson correlation test (*p-value*<0.05 for all comparisons) (Figure S2A). The close agreement between simulated spectra and experimental data confirmed the validity of our simulation protocol. For all backbone atoms, shifts correlation coefficient *R* was more than 0.76. The deviation of HN shifts calculated from simulation structures is higher than in other backbone atoms. This could be explained by the disordered nature of the AF1c and the different geometries of protons interacting with solvent molecules or backbone oxygen atoms. As such, HN shifts are the most sensitive to the protein structure. The root mean square error (RMSE) between experimental and predicted spectra was biggest in case of nitrogen shifts – 1.322 ppm. In other cases, it was lower when 0.5 ppm, all these deviations are within the expected range for the SPARTA+ error ^57^. This result corresponds to the best values reported in the literature for the disordered proteins ^64^. Additionally, we compared secondary NMR shifts, corrected to the random coil components (Figure S3B), which also revealed a significant overlap between simulated and experimental data. However, there were some deviations in the nitrogen shifts, showing some destabilization of the helical conformation of residues 215-226 in comparison to experimental shifts. Another deviation between the experimental and simulation results was observed in the shifts at the C-terminus of the protein for the nitrogen and alpha carbon atoms (Figure S3B). Despite these deviations, the model correctly describes the random nature of the AF1c chain ^64^. The low *p-value* and RMSE between simulated and experimental signals validates our simulation protocols, which we then use to study the phosphovariants, for which NMR data is lacking to date.

### Sequential phosphorylation opens up protein structure

To quantify the effect of phosphorylation on the protein geometry, we visualized distributions of the radius of gyration (*Rg*) and the *SASA* (Figure 2 A,B). The NP molecule has the most compact shape, and upon phosphorylation, AF1c elongates and becomes more solvent-exposed. This trend is expected as more phosphorylation leads to more negative charge in the chain. Analysis reveals that the *Rg* distributions of double and triple phosphorylated variants are relatively close. A single phosphorylation of Ser 211 alters the *SASA* and *Rg* more than a single phosphorylation of Ser 203. The *SASA* metric appears to be more sensitive to the protein structure, and in the case of phosphovariants two-peak distributions are observed, suggesting the co-existence of compact and solvent exposed protein conformations. This indicates that local site-specific changes contribute to regulation of overall compaction.

**Figure 2:**
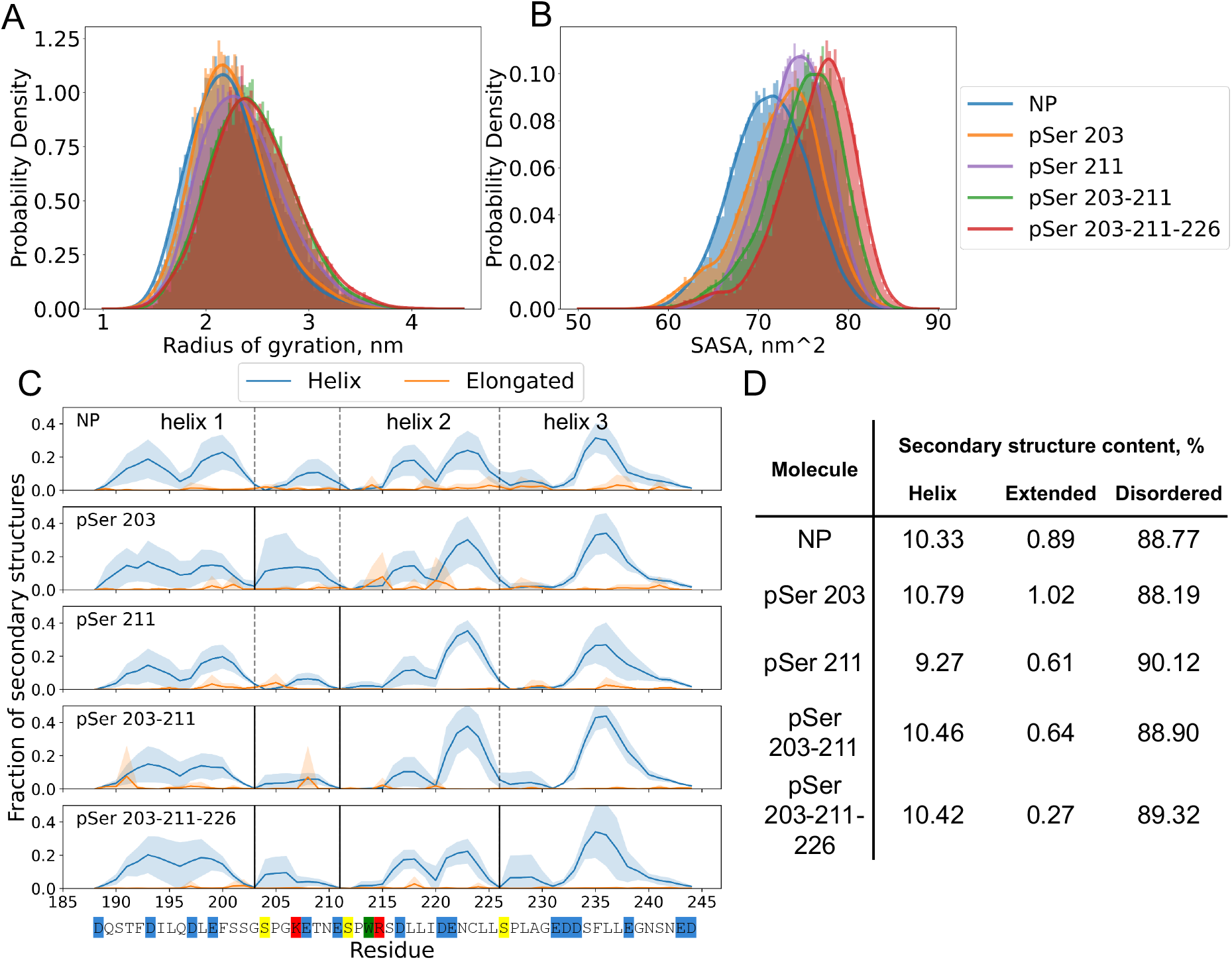
An increase in the protein net charge opens up its structure and promotes the redistribution of secondary structures. The radius of gyration (**A**) and the Solvent Accessible Surface Area (**B**) distributions for different phosphovariants are displayed. (**C**) The fraction of secondary structure per residue for different phosphovariants. Blue – helical structure; orange – extended. Vertical lines representing phosphorylation sites, dashed – non-phosphorylated, solid – phosphorylated. (**D**) The average secondary structure for different phosphovariants. Secondary structure was calculated from combined trajectories for each phosphorylation type with help of DSSP tool ^61^.

The observed radii of gyration are close to one another, and are consistent with theoretical expectations from the theory of disordered proteins ^65^. Our simulations yield an *Rg* for *NP* of 2.2±0.4 nm, which is slightly smaller than what is predicted for excluded volume (scaling exponent 0.6; the bond length 2.2 nm) of the chain, i.e. 2.5 nm. Interestingly, sequential phosphorylation shifts *Rg* to this limit. The pSer 203-211 and pSer 203-211-226 variants yield radii of 2.5±0.4 nm. Such an increase in *Rg* agrees with earlier literature ^65,66^.

### Phosphorylation locally modulates helical structures

Next, we studied the secondary structure content of different phosphovariants, using the DSSP tool ^61^. As shown in Figure 2C the changes in helical structure distribution take place in close proximity of phosphorylation sites. Interestingly, the time-averaged amount of secondary structures was almost identical for all variants, see Figure 2D.

The secondary structure distribution of NP molecule corresponds well to what was expected from previous NMR experiments ^58,67^. More specifically, three helical regions could be observed: 189-203 (helix 1), 215-226 (helix 2), 231-245 (helix 3). The three regions mentioned above appear clearly in the NMR data as regions with relatively high propensity for helix formation^58,67^. Ser-Pro residues separate these helical regions. We note that complete helix formation in each helical region is a rare event, as it could be seen in Figure S7. Instead, we typically see a single helix turn. However, helical structure distribution, observed in simulations of NP phosphovariant, nicely follows secondary structure propensity reported previously in NMR analysis done by *Kim et al* ^58^. Single phosphorylation of the 203 site promotes an increase of the helical conformation in the helix 1 fragment. This could be attributed to charge interaction between phosphoserine and lysine in the sequence: pSer-Pro-Gly-Lys. A proline residue is known to be not only a helix breaker, but also a helix initiator residue. In combination with the small Gly residue, it tends to form a charge interaction between pSer and Lys, which are suitable for hydrogen backbone bonds in a helical structure. In contrast, pSer211 does not stabilize helix 2. In this case, the phosphoserine and the arginine in motif pSer-Pro-Trp-Arg are separated by tryptophan, which competes with Arg in interaction with pSer. This competition makes it harder to create stable hydrogen bonds and to form a helix. The same could be observed in the double phosphorylated variants. The pSer 203-211 and pSer 203-211-226 variants have more stable helix 1 and helix 3, respectively, than other phosphorylation variants. This could be attributed to structure elongation and the increased role of the short-range contacts within the protein backbone.

### Contact analysis reveals distinct subpopulations

Next, we focus on intra-chain contact analysis. We identified important intra-chain contacts by performing a principal component analysis (PCA) to determine and visualize phosphorylation-induced patterns in contact space. First, each frame from the simulation was represented with a vector of protein intra-chain contacts, with a cutoff of 0.5 nm. Then PCA was performed on the trajectories of all phosphovariants projecting them onto the same space. While no single trajectory fully covers the entire PCA space, there is significant overlap between independent replicates of the same phosphovariant. This highlights the necessity of multiple replicates to achieve sufficient sampling. Note that the PCA is an unsupervised method, where the identified components are defined solely by data variability.

The first two eigenvectors of the PCA show the importance of short-range contacts (Figure 3A), identifying mainly the ones that form helical secondary structure. The first principal component could be attributed to the formation of helix 3 and some destabilization of helix 1, while the second principal component represents stabilization of helix 1 and helix 2, see Figure 3A. These eigenvectors represent 3.45% and 3.17% of data variance.

**Figure 3:**
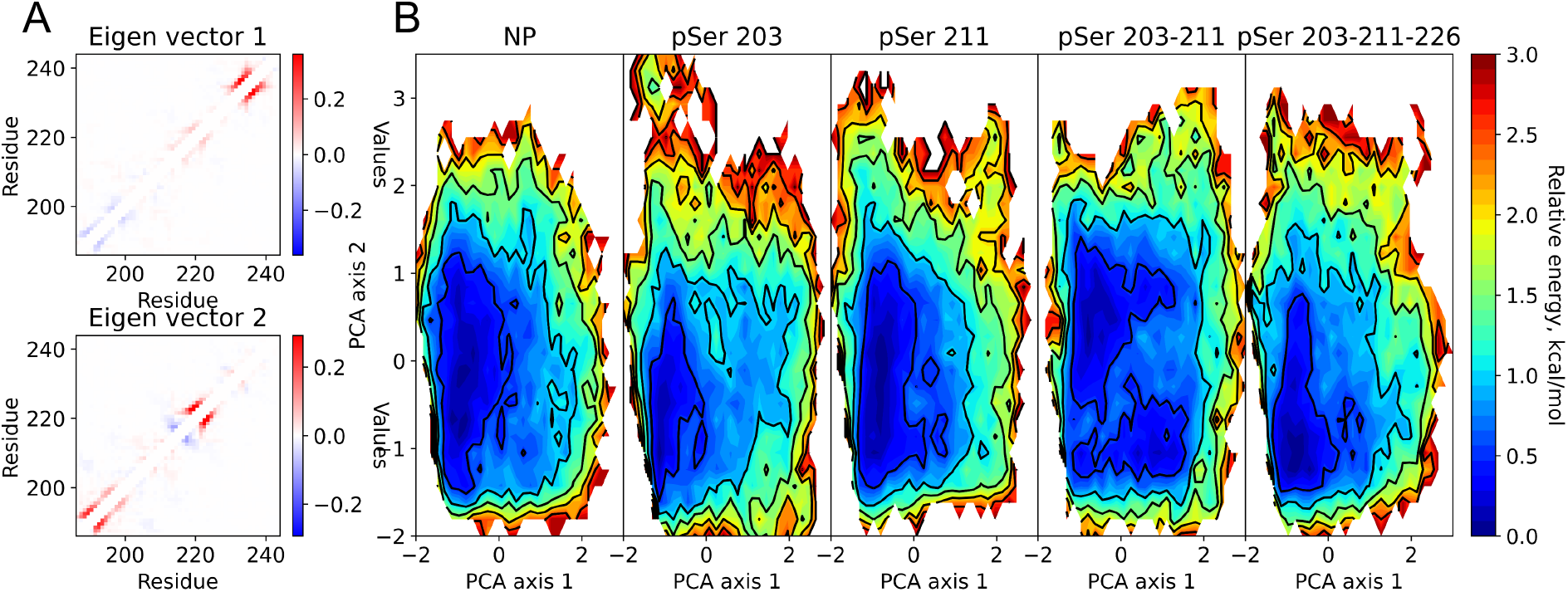
Principal component analysis identifies folding of helices 1 and 2 in AF1c independently from folding of helix 3. (**A**) Contributions to the first two eigenvectors of the PCA transformation per residue. (**B**) Energy landscape of different phosphorylation types.

Our PCA analysis of the studied phosphovariants revealed distinct conformational distributions, which we plotted as energy landscapes in Figure 3B. The energy landscapes of NP and pSer 211 are close to each other: a shallow energy basin is extended along PCA axis 2, showing minima along the two-helix component (eigen vector 2) together with unstructured conformations. The pSer 203 variant contains a more pronounced axis 1 component, and a less pronounced axis 2 character. Importantly, pSer 203-211 has a unique pattern of energy surface, featuring two wells: one representing conformations with stable helices 1 and 2 and the second one representing conformations with stable helix 3. This double phosphovariant has one percent higher amount of secondary structure than others. The most stable conformations of triple phosphovariant are disordered with a well at the bottom-left side of the energy landscape; this makes conformation of the triple variant close to that of NP and pSer 211. The representative clusters for each energy minimum could be found in Figure S12.

### Circuit topology as a tool to identify compact structures

To better resolve transient conformations in the predominantly disordered AF1c structures, we developed and applied a novel circuit topology-based approach (Figure 4A) and classified disordered conformations based on the complexity of their topological structure. The contact analysis presented in the previous section revealed the importance of secondary structures formed by short-range contacts. Topological arrangements of contacts, particularly non-local ones, may also be informative in understanding the conformational dynamics of disordered chains on a more global level. Importantly, AFc1 does not fold in isolation within the timescale of our simulations, consistent with previous experimental reports ^58,68^. However, they may form Disordered Compact States (DCS), which can be resolved using CT.

**Figure 4:**
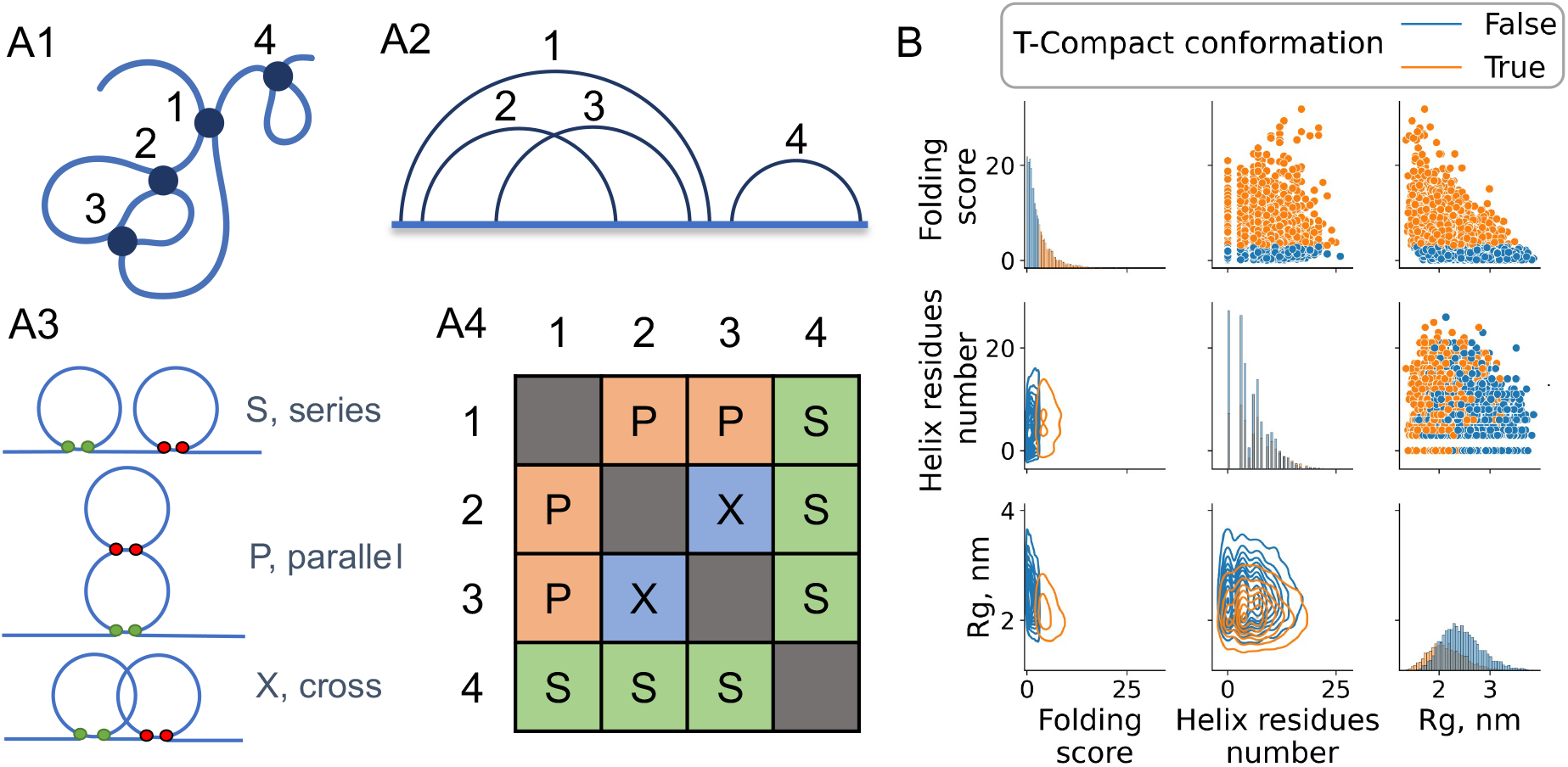
Circuit topology (CT) as metric to track conformational changes in IDPs. (**A1**) CT classifies the relation between intra-chain contacts in a protein. (**A2**) Circuit diagram (arc diagram), where arcs represent contacts between two residues, clearly reveals how contacts and associated loops are entangled. In this example, a circuit is shown which includes four contacts. (**A3**) A given contact pair in circuits could have three basic arrangements: series (S), parallel (P), and cross (X). (**A4**) A protein structure represented with a CT matrix. In the CT matrix, rows/columns represent contacts, and each cell represents the relationship between these contacts. (**B**) The pairwise relationships between the radius of gyration (Rg), helix residues number, and folding score (see Methods for definition) for all phosphovariants combined. On the diagonal are the histograms of Folding Score, amount of helix residues, and Radius of gyration. In the upper triangle of plots grid the pairwise relations are shown as a scatter plots, in lower triangle same data represented in a contour plot representation. Colors represent the distribution of compact conformations (orange) and solvent exposed (blue) conformations. It could be seen, that on average topologically compact conformations have smaller radius of gyration. Decrease in radius shows bigger score.

CT formalizes the pairwise arrangement of contacts in a chain, which can be in Series (S), in Parallel (P), or cross each other (X), as shown in Figure 4A. Over the course of simulations, molecules undergo conformational and topological changes. As could be seen in Figure S4 disordered chains typically have a small number of topological relationships. This could be explained by the low number of contacts that protein has in a solvent exposed – disordered state. However, upon compaction the number of all relations increases. Often, increases in P, S and X relationships correlate, but there are some differences (Figure S4). This observation shows that all three topological relations are important for characterization of folding, but normalization is needed to incorporate them equally. The simplest approach is to use a linear combination of P, S and X with certain weights. For disordered proteins, it is not straight forward to select the most important type or relation; as such, we decided to normalize all three to average observed numbers. This gives the *CT Folding Score* defined as: 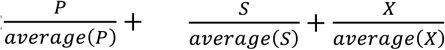 where S is – the number of series relations, P - is the number of parallel relations, X - is the number of cross relations between the contacts; each is normalized on average number of relations observed for all snapshots of the molecule. A conformation is defined as “topologically compact” (T-compact) if it has a *folding score* more than the average folding score, which for all cases is equal to 3. In other words, if the amount of each of relations is more than average, then the structure is called T-compact (Figure S5). Considering that the average state for a disordered protein is “disordered”, one can analyze deviation from this average which signifies the emergence of T-compact states. Such an approach helped us to efficiently distinguish topologically compact and solvent exposed conformations (Figure 5). Formulated in such a way, *CT folding score* gives nice contrast during compaction events as could be seen in Figure S6. In this figure, compaction event corresponds to decrease in Rg and SASA, and significant increase in *CT Folding Score*. Compared to simply counting the overall number of contacts, this approach is more sensitive to the formation of non-local contacts and contains information about the arrangement of contacts. Consequently, the protein conformations classified as a “compact”, using *CT Folding Score*, have a smaller radius of gyration *Rg*, despite a varying amount of helical structure (Figure 4B). Another advantage of CT Folding Score is that it fluctuates less than geometrical metrics. With stable contacts formation the *Folding Score* would not fluctuate, while radius of gyration and SASA could change because of movements of solvent exposed loops (Figure S6, Figure 4B). As such, topologically compact states may occasionally exhibit high Rg or SASA and therefore may not be “geometrically” compact (G-compact).

**Figure 5:**
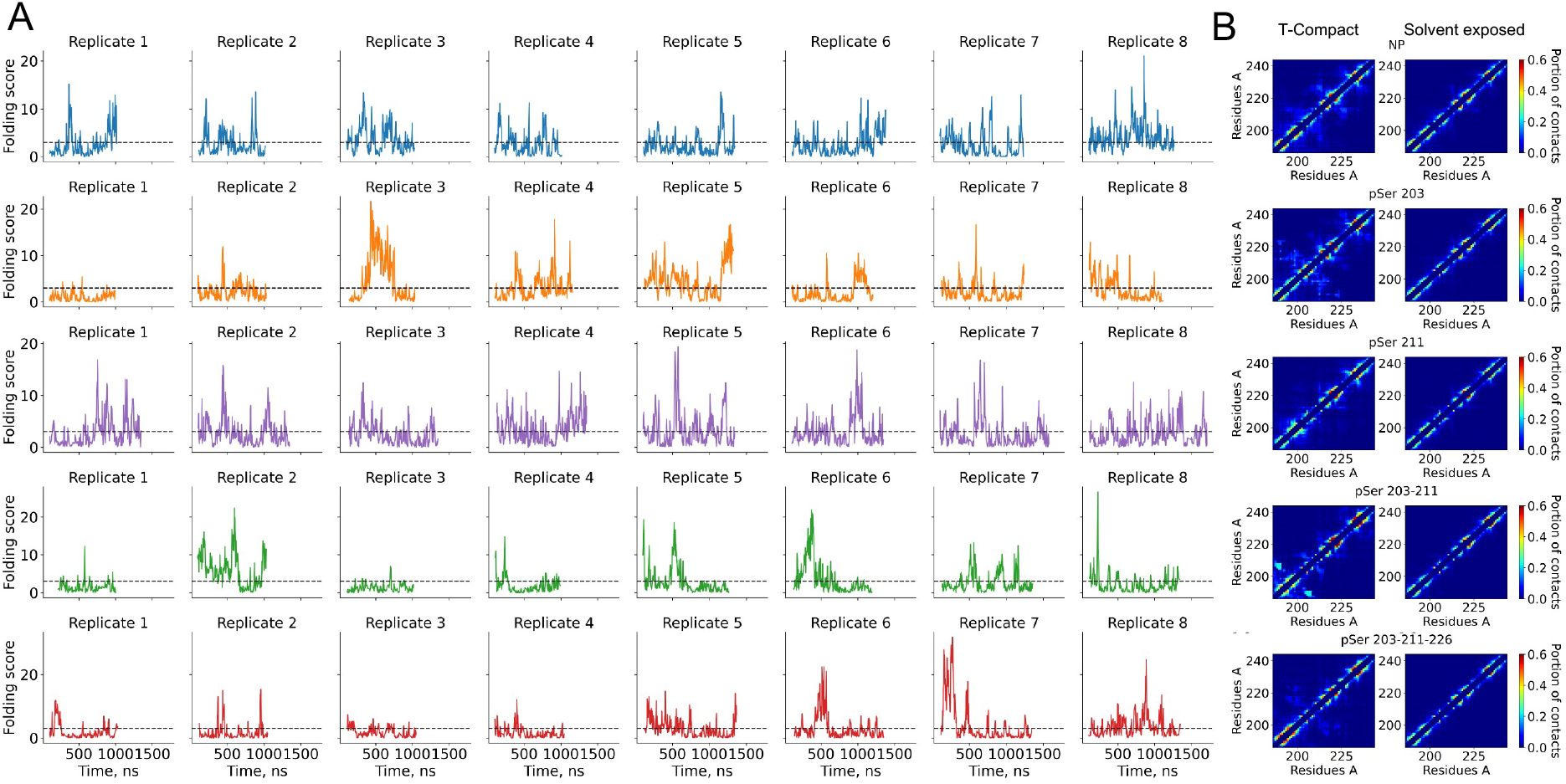
Circuit topology identifies long-range contacts in disordered protein conformations. (**A**) CT Folding score dynamics during the simulation for different phosphorylation variants. The dashed horizontal line shows the threshold that differentiates solvent exposed and topologically compact structures. (**B**) The averaged contact matrix for compact and solvent exposed protein conformations.

Alternative threshold could be suggested for folding simulations, such that threshold would separate disordered and completely folded structures. This could be done after obtaining histogram of the score and positioning threshold between these two conformation classes, at lowest probability point, that represents transition state.

The phosphorylation of AF1c affects the propensity and kinetics to form disordered compact conformations in a site-specific manner. In Figure 5A, the *CT Folding Score* dynamics for each phosphovariant and corresponding independent trajectories are represented. Considering *CT Folding Score* as a metric of protein compaction, a burst can be interpreted as a compaction event. In the case of pSer 203, pSer 203-211, and pSer 203-211-226 molecules, several long living compact topologies were observed. Interestingly both NP and pSer 211 have kinetics patterns that are distinct from those of other phosphovariants. This difference could be explained by fast transitions between disordered and helix containing states, consistent with the absence of a free energy barrier between them (see Figure 3). In the case of double and triple phosphovariant, stretches of a low score followed by long-living bursts were observed, reflecting periods of extended protein structures, followed by sudden compaction.

Contact maps show that solvent-exposed disordered structures are similar between all phosphovariants of AF1c. With the *CT Folding Score*, we can now analyze the effect of phosphorylation on contact formation propensity separately for disordered compact and disordered solvent-exposed conformations (Figure 5B). The contact maps show that the topologically compact conformations contain not only non-local contacts, but also have a higher propensity to form helical structures. Compact conformations of pSer 203 and pSer 203-211-226 molecules show interactions between helix 1 and helix 2. Interestingly, in the case of pSer 203, residues 203-211 form a helix (Figure 3B), but in pSer 203-211, a disordered region could be observed between helix 1 and residues 203-211. Some helices contain kinks within them (Figure 5B). When the helix is interrupted by a disordered region, we observed that the two helix parts tend to interact with each other. Interestingly, despite NP having the smallest average *Rg* and *SASA*, its compact shape is formed by intra-helical region contacts and increased non-ordered contacts in the 203-211 region. Although the solvent-exposed conformations still contain a significant amount of helical structures, they lack non-local contacts.

Same trends could be observed by analyzing the topology matrices (Figure S10). Different phosphovariant shows different propensity for long non-local contacts formation. Parallel types of topological relations is the most sensitive for such contacts. The pSer 203 and pSer 203-211 phosphovariants have distinguished contacts between non-helical region and helix 2 (residues 210-220). The average amount of topological relationships decreases upon phosphorylation, which correlates with increases in Rg and SASA. Interestingly, phosphorylation of Ser 203 increases the number of parallel relations between helix 1 and helix 2. Some interaction could be detected already in NP case, but phosphorylation of 203 increases such interactions, while single phosphorylation of Ser 211 decreases it.

### The conformational kinetics is modulated by phosphorylation

Finally, we characterized the dynamics of phosphovariant with different folding metrics. We have done autocorrelation analysis of *CT Folding Score*, radius of gyration, SASA, and amount of helical structure (Figure 6A, Figure S9A). In all these metrics, autocorrelation of NP and pSer 203, and double and triple phosphorylated AF1c forms two groups. First with slow dynamics, with time of half autocorrelation decay of 30 ns, and fast dynamics with time of half decay of 10 ns. Interestingly, Rg and SASA autocorrelation of AF1c pSer203 lays between these two groups, but *CT Folding Score* and number of helix residues goes to fast dynamics group. The slow dynamics for NP could be explained by more compact solvent exposed structure (Figure 2A,B). In this case, significant structural fluctuations occur on slower timescales. In the case of AF1c pSer 203, slow autocorrelation could be explained by the formation of the disordered compact states. The most prominent examples are replicates 3 and 5. During the simulation of AF1c, pSer 203-211 and pSer 203-211-226 topologically compact states were formed as well, such as in replicate 7 triple phosphorylated variant. However, the dominant state was solvent exposed, which dominated the autocorrelation function. Interestingly, no compact disordered states were observed for AF1c pSer 211. Autocorrelation functions of individual trajectories can be found in Figure S9B.

**Figure 6:**
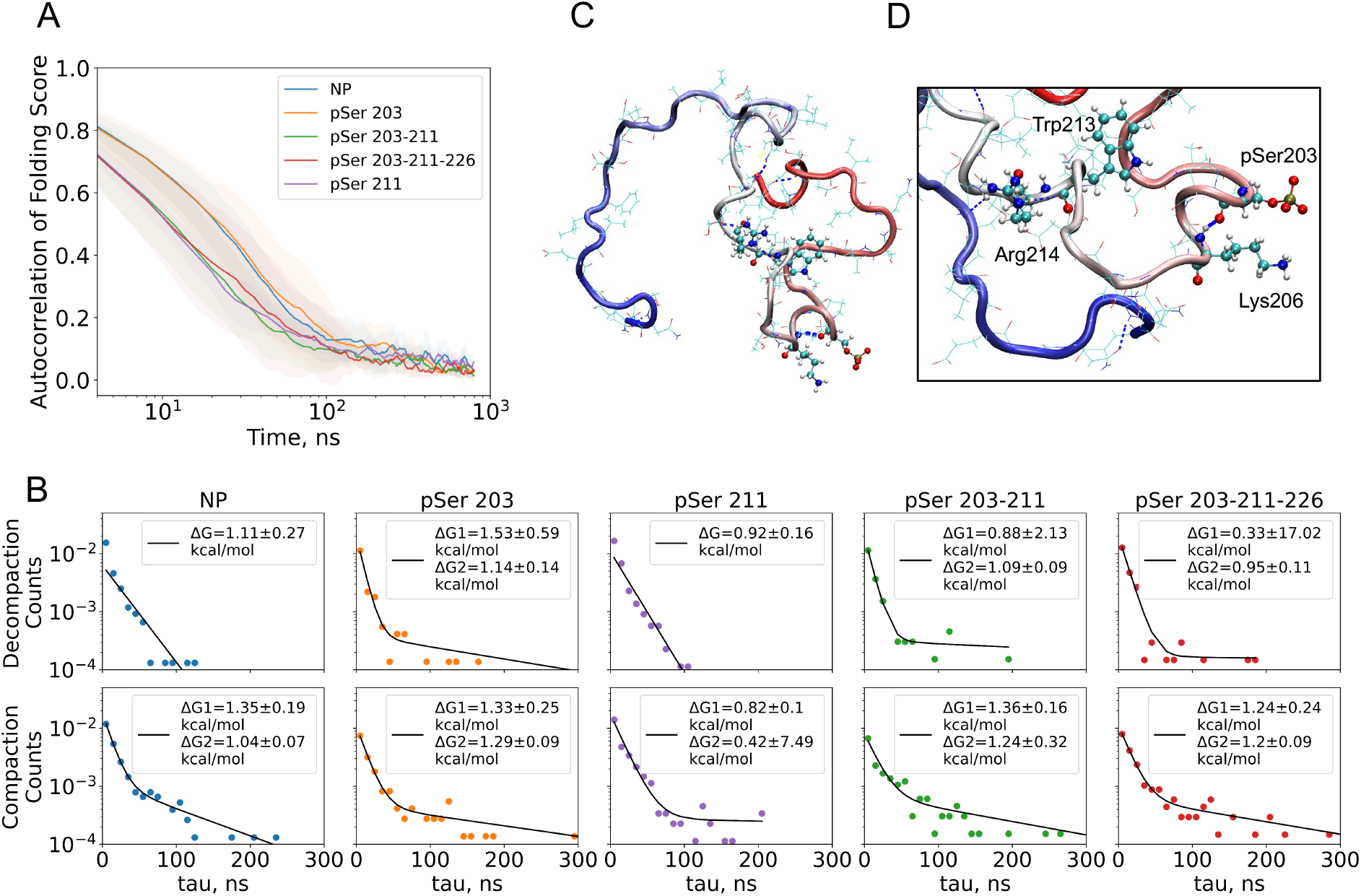
Phosphorylation promotes formation of long-living compact states and alters distribution of protein structures. (**A**) Autocorrelation of CT Folding Score for different phosphovariants. NP and S211P show compaction events with 10-30 ns lifetimes, while S203P, S203-211P and S203-211-226P show relatively slow (50 – 200 ns) compaction events. (**B**) Dwell-time histograms fitted with a single and double exponential decay function. The histograms were constructed from the times protein was in the disordered compact state (Decompaction) and solvent exposed state (Compaction), then counts were divided on the simulation time for the activation energy ΔG calculation. Here, we presented the histogram fits with single and double exponent function; double exponent fits were done on Decompaction times for pSer203, pSer 203-211, and pSer 203-211-226. Slow exponent represents the dynamics of compact disordered states. These events are rare and have long durations, that characterize the form of the curve. (**C**) Disordered compact shape of S203P at frame 450 (CT Folding Score peak). Colors show the amino acid number: from red – N-terminal region to blue – C-terminal region. In this image, one can readily see that most of the protein is in direct contact with solvent, while there is a contact between N-terminal region and the middle of the protein (compare with contact map for S203P/compact in Figure 5B). (**D**) Close look at the helix formed by interaction of pSer203-Lys206. This helix is stabilized by charge interaction and backbone hydrogen bond formation (blue dashed line)

The dwell time distribution of the *CT Folding Score* showed phosphorylation-induced stabilization of disordered compact states. In contrast to autocorrelation analysis, dwell analysis is more sensitive to *rare events*. Folding score traces were transformed to dwell time histograms by applying a threshold that represents the topologically compact state. Analysis of dwell time revealed a long-living component for decompaction of pSer 203, pSer 203-211 and pSer 203-211-226 molecules (Figure 6B), suggesting the stabilization of compact conformation in these cases. For all phosphovariants slow compaction was observed. These results suggest that compaction events are phosphorylation independent, but decompaction and compact state stabilization is regulated by phosphorylation. In other words, topologically contact states are stabilized by interactions with phosphoserines (Figure 6C, D). The partial screening of the positively charged Lys 206 (in case of AF1c pSer 203,) and Arg 214 (AF1c pSer 203-211) caused by phosphorylation promotes change in the average contact map of the protein compact state (Figure 5B). After “neutralization” of charge of these residues, their hydrophobic properties remain, and they could be engaged in hydrophobic contacts, that stabilizes disordered compact states. The energy barrier in non-phosphorylated and pSer 211 cases could be easily crossed, but lowering of the ion concentration slows down the compaction kinetics and promotes the formation of the long living disordered compact states, in the same manner as in the case of multiple phosphorylations at native salt conditions (Figure S13C, D). All these observations show the role of the electrostatic interaction balance on the disordered protein conformational ensemble and kinetics.

## Discussion & Conclusions

By performing more than 40 microseconds of atomistic unbiased MD simulations, this study demonstrates that the AF1c subunit of the GR and its phosphovariants are disordered and that site-specific phosphorylation modulates the kinetics and the probability of forming disordered compact states. NP and pSer 211 variants show many short-living compaction events. This could be attributed to the formation of weak contacts between distinct protein regions forming extended (in terms of DSSP classification) secondary structure (Figure S7A, C). In the case of pSer 203 and pSer 203-211 variants, we could observe stable extended structure contacts more often, and these contacts are responsible for the majority of the compact, yet disordered states. There are interesting exceptions, however, such as a long living disordered folded state in pSer 203 observed in replicate 3 (Figure S7B, Figure 6 B C) and in pSer 203-211 replicate 6 (Figure S7D). These states cannot be attributed to the extended structure formation, but have a higher content of helical structure, including helix in 203-211 region in case of pSer 203. Solvent-exposed conformations are rarely followed by disordered compact conformations, stabilized by extended structure or helix 1 and helix 2 interactions (Figure S7). The triple phosphorylated variant pSer 203-211-226 has a high propensity for helical structure formation (Figure 2C), and pSer 203-211-226 has the most solvent-exposed structure (Figure 2AB). Such a combination could explain the dynamics of this variant’s *CT Folding Score*.

Experimental research has suggested that phosphorylation of Ser211 residue promotes folding of AF1c ^17, 18, 25^, which likely happens within timescales much longer (dozens of microseconds) than what our study probes. However, we have made observations that are in line with phosphorylation-induced folding (as observed in previous CD analysis ^18, 25^). For example, we have observed helix formation of the 203-211 fragment after phosphorylation of Ser203 residue (Figure S7B, Figure 6D). At low salt conditions (25 mM NaCl, pSer 211) (Figure S13A, I), pSer211-Pro212-Trp213-Arg214 formed stable helices in our simulations, in the same manner as observed earlier for pSer203-Pro204-Gly205-Lys206. These observations support the previous suggestion that pSer-Pro tends to be a helix initiator motif ^69^. Thus, phosphorylation may be associated with stabilization of helical conformations by charge interaction and hydrogen bonds. These helix formation seeds could grow into well-defined helical structures and eventually stabilize folds on long time scales.

The reversible phosphorylation of GR plays a key role in modulating the GR response by influencing its cellular localization, and it is regulated by the balance between the phosphorylating targeted kinases and dephosphorylating phosphatases that will interact with the receptor ^2, 70^. Our study reveals the detailed effect of phosphorylation on the conformational diversity of AF1c, suggesting that GR adopts different topological dynamic states as it translocates from the cytoplasmic to the nuclear environment. This sheds new light on how phosphorylation of Ser203 and Ser211 results in enhanced nuclear localization and subsequent increase of GR transcriptional activity as previously reported ^2, 22^. The ability of AF1c phosphovariants to adopt multiple structural changes likely creates distinct binding surfaces within the NTD, enabling GR to selectively interact with different regulatory complexes along its signal transduction pathway. Interestingly, we have observed that pSer 203-211 has the highest propensity for secondary structure formation. This phosphovariant is localized in the perinuclear region and is believed to be the precursor of the active form of GR (pSer211), found in the nucleus ^21^. In this context, it is noteworthy that pSer 211 and pSer 203-211 exhibit distinct structural propensities, indicating that different conformations may be needed to transition between the two phosphorylated states. Whereas Ser 203 can be phosphorylated by both cyclin-dependent kinases (CDKs) and ERK kinases, p28 MAPK is the one that fully activates GR by phosphorylating Ser211 after ligand-binding, suggesting that these different structural states can influence the interaction of GR with different kinases ^22^. On the other hand, the shift from one state to another is significantly influenced by the presence of phosphatases interacting with GR and reverting its modifications, which are also involved in the nuclear import of the ligand-activated GR ^22^. Our results are in line with the experiments performed by Garza et al. 2010, who demonstrated that phosphorylation at Ser211 leads to significant structural changes in the GR AF1 domain, supporting the hypothesis that this modification induces a more ordered and functionally relevant conformation capable of interacting with several transcriptional coregulators ^25^. We hypothesize that each phosphorylation-driven conformational state could favor the interaction of GR with specific partners by altering the accessibility of AF1c binding sites, thereby modulating its activity and transcriptional outcomes in a context-specific manner.

GR may be phosphorylated in more ways than this research covers. Zhen Wang *et al*. ^23^ reported observation of pSer211-pSer226, pSer203-pS226, and pSer226 phosphorylation variants. The AF1c pSer203-pS226 is characteristic of the perinuclear region. Of particular interest is the comparison of nucleus-localized AF1c pSer211, AF1c pSer226, and AF1c pS211-pSer226, which may have different transactivation activity. The listed combinations of phosphorylation could exhibit unique propensities of disordered compact state formation, which may be linked to their biological role. Combining all-atom MD simulations with CT analysis can readily be used to analyze these variants, which we plan for the follow up research.

The central result of this research is a new methodology for IDP analysis, which we demonstrated using the GR AF1 analysis as a proof-of-concept. The *CT Folding Score*, a circuit topology-based metric, calculates the normalized number of series, parallel and cross types of contact relations. Using *CT Folding Score*, we quantified the probability of disordered compact state formation, where the protein chain is solvent-exposed, yet certain residues form transient (non-local) contacts. In contrast to CT Folding Score analysis, characterization of such states by secondary structure content, *Rg* or *SASA* is not productive due to large structural variations and the incompleteness of these representations (Figure S6). However, compact disordered states could be efficiently defined using the language of circuits. Figure S8 shows that protein conformations with higher *CT Folding Score* usually have a smaller Rg and SASA, independent from helical structure. The stable contact formation is the essence of protein folding. Since *CT Folding Score* contains information about protein chain entanglement, it can sensitively detect protein compaction and even folding events. One can consider utilizing the *CT Folding Score* as a coordinate of folding. In future applications this metric could be used in meta-dynamics folding simulations ^71^. Depending on the application, users may refine the *CT Folding Score*, for example, by introducing different weights or normalizing it by maximum topological relations. In this research the threshold was optimized to track the deviation of topology from the main – disordered state. In other cases, the cut-off could be adjusted. For example, for folding simulations it is advised to use the transitions state topology for defining the threshold between solvent exposed and folded states.

Fully or partly intrinsically disordered proteins account for a significant fraction of human proteome, yet our understanding of their conformational regulation is limited. It is known that IDPs often get stabilized by binding to their interacting partners, but this is not the only way that disordered to ordered transitions happen. Disordered compact states can be seen in isolated IDPs depending on their sequence, post-translational modification, and solvent condition ^4,72^. In contrast to the molten globule state, commonly seen as a transition state in the folding of stably folded proteins, disordered compact conformations (which we may also refer to as “disordered connected” conformations) do not contain a lot of secondary structure. In this way, they could be considered as precursors of a molten globule state. Despite structural instability, formation of compact disordered states is linked to function ^5,73^. Importantly, reversible protein phosphorylation provides a major regulatory mechanism in conformational dynamics and compaction of IDPs ^16,74^, which can be effectively characterized using the topology-based approach developed in this study. Chains that form transient contacts at various time scales and with different interconnected chain regions can form “circuits” that feature both disorder and compactness simultaneously.

## Author Contributions

Conceptualization & Project Administration: A.M. Investigation: V.A. Formal Analysis: V.A. Software: V.A. Methodology: V.A., A.M. Visualization: V.A., J.v.N., A.M. Writing & Draft Preparation: V.A., A.M. Writing-Review & Editing: V.A., A.J.P., E.E., J.v.N., A.M.

## Note

The authors declare no conflict of interest.

## Acknowledgements

The authors would like to thank the technical support of Vahid Sheikhhassani from Medical Systems Biophysics and Bioengineering Laboratory, Leiden University. This work was performed using the Academic Leiden Interdisciplinary Cluster Environment (ALICE), as well as Snellius, the Dutch National supercomputer hosted at SURF (grant number: EINF-7466).

## Supporting Information Available

The Supporting Information is available free of charge on the ACS Publications Web site.

The circuit topology-based fold analysis tool developed in this study is available at: https://github.com/circuittopology/CT_Folding_Score

The simulation trajectories have been deposited in Zenodo repositories and can be found at: Doi: 10.5281/zenodo.13820169

Doi: 10.5281/zenodo.13822438

## References

(1) Uversky, V. N.; Dunker, A. K. Understanding protein non-folding. Biochim Biophys Acta 2010, 1804 (6), 1231–1264. DOI: 10.1016/j.bbapap.2010.01.017.

(2) van der Lee, R.; Buljan, M.; Lang, B.; Weatheritt, R. J.; Daughdrill, G. W.; Dunker, A. K.; Fuxreiter, M.; Gough, J.; Gsponer, J.; Jones, D. T.; et al. Classification of intrinsically disordered regions and proteins. Chem Rev 2014, 114 (13), 6589–6631. DOI: 10.1021/cr400525m.

(3) Chowdhury, A.; Nettels, D.; Schuler, B. Interaction Dynamics of Intrinsically Disordered Proteins from Single-Molecule Spectroscopy. Annu Rev Biophys 2023, 52, 433–462. DOI: 10.1146/annurev-biophys-101122-071930.

(4) Das, R. K.; Pappu, R. V. Conformations of intrinsically disordered proteins are influenced by linear sequence distributions of oppositely charged residues. Proc Natl Acad Sci U S A 2013, 110 (33), 13392–13397. DOI: 10.1073/pnas.1304749110.

(5) Wang, D.; Wu, S. W.; Wang, D. D.; Song, X. Y.; Yang, M. H.; Zhang, W. L.; Huang, S. H.; Weng, J. W.; Liu, Z. J.; Wang, W. N. The importance of the compact disordered state in the fuzzy interactions between intrinsically disordered proteins. Chem Sci 2022, 13 (8), 2363–2377. DOI: 10.1039/d1sc06825c.

(6) Tompa, P.; Schad, E.; Tantos, A.; Kalmar, L. Intrinsically disordered proteins: emerging interaction specialists. Curr Opin Struct Biol 2015, 35, 49–59. DOI: 10.1016/j.sbi.2015.08.009.

(7) Golovnev, A.; Mashaghi, A. Generalized Circuit Topology of Folded Linear Chains. iScience 2020, 23 (9), 101492. DOI: 10.1016/j.isci.2020.101492.

(8) Scalvini, B.; Sheikhhassani, V.; Mashaghi, A. Topological principles of protein folding. Phys Chem Chem Phys 2021, 23 (37), 21316–21328. DOI: 10.1039/d1cp03390e.

(9) Schullian, O.; Woodard, J.; Tirandaz, A.; Mashaghi, A. A Circuit Topology Approach to Categorizing Changes in Biomolecular Structure. Front Phys-Lausanne 2020, 8. DOI: ARTN 5 10.3389/fphy.2020.00005.

(10) Heidari, M.; Schiessel, H.; Mashaghi, A. Circuit Topology Analysis of Polymer Folding Reactions. ACS Cent Sci 2020, 6 (6), 839–847. DOI: 10.1021/acscentsci.0c00308.

(11) Scalvini, B.; Sheikhhassani, V.; van de Brug, N.; Heling, L.; Schmit, J. D.; Mashaghi, A. Circuit Topology Approach for the Comparative Analysis of Intrinsically Disordered Proteins. J Chem Inf Model 2023, 63 (8), 2586–2602. DOI: 10.1021/acs.jcim.3c00391.

(12) Kadmiel, M.; Cidlowski, J. A. Glucocorticoid receptor signaling in health and disease. Trends Pharmacol Sci 2013, 34 (9), 518–530. DOI: 10.1016/j.tips.2013.07.003.

(13) Dieken, E. S.; Miesfeld, R. L. Transcriptional transactivation functions localized to the glucocorticoid receptor N terminus are necessary for steroid induction of lymphocyte apoptosis. Mol Cell Biol 1992, 12 (2), 589–597. DOI: 10.1128/mcb.12.2.589-597.1992.

(14) Dahlman-Wright, K.; Almlof, T.; McEwan, I. J.; Gustafsson, J. A.; Wright, A. P. Delineation of a small region within the major transactivation domain of the human glucocorticoid receptor that mediates transactivation of gene expression. Proc Natl Acad Sci U S A 1994, 91 (5), 1619–1623. DOI: 10.1073/pnas.91.5.1619.

(15) Anbalagan, M.; Huderson, B.; Murphy, L.; Rowan, B. G. Post-translational modifications of nuclear receptors and human disease. Nucl Recept Signal 2012, 10, e001. DOI: 10.1621/nrs.10001.

(16) Weikum, E. R.; Knuesel, M. T.; Ortlund, E. A.; Yamamoto, K. R. Glucocorticoid receptor control of transcription: precision and plasticity via allostery. Nat Rev Mol Cell Biol 2017, 18 (3), 159–174. DOI: 10.1038/nrm.2016.152.

(17) Kumar, R.; Thompson, E. B. Role of Phosphorylation in the Modulation of the Glucocorticoid Receptor’s Intrinsically Disordered Domain. Biomolecules 2019, 9 (3), 95. DOI: 10.3390/biom9030095.

(18) Khan, S. H.; McLaughlin, W. A.; Kumar, R. Site-specific phosphorylation regulates the structure and function of an intrinsically disordered domain of the glucocorticoid receptor. Sci Rep 2017, 7 (1), 15440. DOI: 10.1038/s41598-017-15549-5.

(19) Almlof, T.; Wright, A. P.; Gustafsson, J. A. Role of acidic and phosphorylated residues in gene activation by the glucocorticoid receptor. J Biol Chem 1995, 270 (29), 17535–17540. DOI: 10.1074/jbc.270.29.17535.

(20) Wang, Z.; Frederick, J.; Garabedian, M. J. Deciphering the phosphorylation “code” of the glucocorticoid receptor in vivo. J Biol Chem 2002, 277 (29), 26573–26580. DOI: 10.1074/jbc.M110530200.

(21) Ismaili, N.; Garabedian, M. J. Modulation of glucocorticoid receptor function via phosphorylation. Ann N Y Acad Sci 2004, 1024, 86–101. DOI: 10.1196/annals.1321.007.

(22) Miller, A. L.; Webb, M. S.; Copik, A. J.; Wang, Y.; Johnson, B. H.; Kumar, R.; Thompson, E. B. p38 Mitogen-activated protein kinase (MAPK) is a key mediator in glucocorticoid-induced apoptosis of lymphoid cells: correlation between p38 MAPK activation and site-specific phosphorylation of the human glucocorticoid receptor at serine 211. Mol Endocrinol 2005, 19 (6), 1569–1583. DOI: 10.1210/me.2004-0528.

(23) Wang, Z.; Chen, W.; Kono, E.; Dang, T.; Garabedian, M. J. Modulation of glucocorticoid receptor phosphorylation and transcriptional activity by a C-terminal-associated protein phosphatase. Mol Endocrinol 2007, 21 (3), 625–634. DOI: 10.1210/me.2005-0338.

(24) Chen, W.; Dang, T.; Blind, R. D.; Wang, Z.; Cavasotto, C. N.; Hittelman, A. B.; Rogatsky, I.; Logan, S. K.; Garabedian, M. J. Glucocorticoid receptor phosphorylation differentially affects target gene expression. Mol Endocrinol 2008, 22 (8), 1754–1766. DOI: 10.1210/me.2007-0219.

(25) Garza, A. M.; Khan, S. H.; Kumar, R. Site-specific phosphorylation induces functionally active conformation in the intrinsically disordered N-terminal activation function (AF1) domain of the glucocorticoid receptor. Mol Cell Biol 2010, 30 (1), 220–230. DOI: 10.1128/MCB.00552-09.

(26) Ponce-Lina, R.; Serafin, N.; Carranza, M.; Aramburo, C.; Prado-Alcala, R. A.; Luna, M.; Quirarte, G. L. Differential Phosphorylation of the Glucocorticoid Receptor in Hippocampal Subregions Induced by Contextual Fear Conditioning Training. Front Behav Neurosci 2020, 14, 12. DOI: 10.3389/fnbeh.2020.00012.

(27) Copik, A. J.; Webb, M. S.; Miller, A. L.; Wang, Y.; Kumar, R.; Thompson, E. B. Activation function 1 of glucocorticoid receptor binds TATA-binding protein in vitro and in vivo. Mol Endocrinol 2006, 20 (6), 1218–1230. DOI: 10.1210/me.2005-0257.

(28) Kumar, R.; McEwan, I. J. Allosteric modulators of steroid hormone receptors: structural dynamics and gene regulation. Endocr Rev 2012, 33 (2), 271–299. DOI: 10.1210/er.2011-1033.

(29) Garza, A. S.; Khan, S. H.; Moure, C. M.; Edwards, D. P.; Kumar, R. Binding-folding induced regulation of AF1 transactivation domain of the glucocorticoid receptor by a cofactor that binds to its DNA binding domain. PLoS One 2011, 6 (10), e25875. DOI: 10.1371/journal.pone.0025875.

(30) Khan, S. H.; Ling, J.; Kumar, R. TBP binding-induced folding of the glucocorticoid receptor AF1 domain facilitates its interaction with steroid receptor coactivator-1. PLoS One 2011, 6 (7), e21939. DOI: 10.1371/journal.pone.0021939.

(31) Mirdita, M.; Schutze, K.; Moriwaki, Y.; Heo, L.; Ovchinnikov, S.; Steinegger, M. ColabFold: making protein folding accessible to all. Nat Methods 2022, 19 (6), 679–682. DOI: 10.1038/s41592-022-01488-1.

(32) Jumper, J.; Evans, R.; Pritzel, A.; Green, T.; Figurnov, M.; Ronneberger, O.; Tunyasuvunakool, K.; Bates, R.; Zidek, A.; Potapenko, A.; et al. Highly accurate protein structure prediction with AlphaFold. Nature 2021, 596 (7873), 583–589. DOI: 10.1038/s41586-021-03819-2.

(33) Evans, R.; O’Neill, M.; Pritzel, A.; Antropova, N.; Senior, A.; Green, T.; Žídek, A.; Bates, R.; Blackwell, S.; Yim, J.; et al. Protein complex prediction with AlphaFold-Multimer. 2022. DOI: 10.1101/2021.10.04.463034.

(34) David Gfeller, O. M. a. V. Z. Expanding molecular modeling and design tools to non-natural sidechains. Journal of Computational Chemistry 2012, 33, 1525–1535. DOI: 10.1002/jcc.22982.

(35) Gfeller, D.; Michielin, O.; Zoete, V. SwissSidechain: a molecular and structural database of non-natural sidechains. Nucleic Acids Res 2013, 41 (Database issue), D327–332. DOI: 10.1093/nar/gks991.

(36) Schrodinger, LLC. The AxPyMOL Molecular Graphics Plugin for Microsoft PowerPoint, Version 1.8. 2015.

(37) Schrodinger, LLC. The JyMOL Molecular Graphics Development Component, Version 1.8. 2015.

(38) Schrodinger, LLC. The PyMOL Molecular Graphics System, Version 1.8. 2015.

(39) Abraham, M. J.; Murtola, T.; Schulz, R.; Páll, S.; Smith, J. C.; Hess, B.; Lindahl, E. GROMACS: High performance molecular simulations through multi-level parallelism from laptops to supercomputers. SoftwareX 2015, 1-2, 19–25. DOI: 10.1016/j.softx.2015.06.001.

(40) Bonomi, M.; Bussi, G.; Camilloni, C.; Tribello, G. A.; Banáš, P.; Barducci, A.; Bernetti, M.; Bolhuis, P. G.; Bottaro, S.; Branduardi, D.; et al. Promoting transparency and reproducibility in enhanced molecular simulations. Nature Methods 2019, 16 (8), 670–673. DOI: 10.1038/s41592-019-0506-8.

(41) Tribello, G. A.; Bonomi, M.; Branduardi, D.; Camilloni, C.; Bussi, G. PLUMED 2: New feathers for an old bird. Computer Physics Communications 2014, 185 (2), 604–613. DOI: 10.1016/j.cpc.2013.09.018.

(42) Robustelli, P.; Piana, S.; Shaw, D. E. Developing a molecular dynamics force field for both folded and disordered protein states. Proc Natl Acad Sci U S A 2018, 115 (21), E4758–E4766. DOI: 10.1073/pnas.1800690115.

(43) Piana, S.; Robustelli, P.; Tan, D.; Chen, S.; Shaw, D. E. Development of a Force Field for the Simulation of Single-Chain Proteins and Protein-Protein Complexes. J Chem Theory Comput 2020, 16 (4), 2494–2507. DOI: 10.1021/acs.jctc.9b00251.

(44) Steinbrecher, T.; Latzer, J.; Case, D. A. Revised AMBER parameters for bioorganic phosphates. J Chem Theory Comput 2012, 8 (11), 4405–4412. DOI: 10.1021/ct300613v.

(45) Hess, B.; Bekker, H.; Berendsen, H. J. C.; Fraaije, J. G. E. M. LINCS: A linear constraint solver for molecular simulations. Journal of Computational Chemistry 1997, 18 (12), 1463–1472. DOI: 10.1002/(sici)1096-987x(199709)18:12<1463::Aid-jcc4>3.0.Co;2-h.

(46) Darden, T.; York, D.; Pedersen, L. Particle mesh Ewald: An N·log(N) method for Ewald sums in large systems. The Journal of Chemical Physics 1993, 98 (12), 10089–10092. DOI: 10.1063/1.464397.

(47) Virtanen, P.; Gommers, R.; Oliphant, T. E.; Haberland, M.; Reddy, T.; Cournapeau, D.; Burovski, E.; Peterson, P.; Weckesser, W.; Bright, J.; et al. SciPy 1.0: fundamental algorithms for scientific computing in Python. Nat Methods 2020, 17 (3), 261–272. DOI: 10.1038/s41592-019-0686-2.

(48) Hunter, J. D. Matplotlib: A 2D graphics environment. Comput Sci Eng 2007, 9 (3), 90–95. DOI: Doi 10.1109/Mcse.2007.55.

(49) Harris, C. R.; Millman, K. J.; van der Walt, S. J.; Gommers, R.; Virtanen, P.; Cournapeau, D.; Wieser, E.; Taylor, J.; Berg, S.; Smith, N. J.; et al. Array programming with NumPy. Nature 2020, 585 (7825), 357–362. DOI: 10.1038/s41586-020-2649-2.

(50) McGibbon, R. T.; Beauchamp, K. A.; Harrigan, M. P.; Klein, C.; Swails, J. M.; Hernandez, C. X.; Schwantes, C. R.; Wang, L. P.; Lane, T. J.; Pande, V. S. MDTraj: A Modern Open Library for the Analysis of Molecular Dynamics Trajectories. Biophys J 2015, 109 (8), 1528–1532. DOI: 10.1016/j.bpj.2015.08.015.

(51) Alibay, I.; Barnoud, J.; Beckstein, O.; Gowers, R. J.; Loche, P. R.; MacDermott-Opeskin, H.; Matta, M.; Naughton, F. B.; Reddy, T.; Wang, L. Building a community-driven ecosystem for fast, reproducible, and reusable molecular simulation analysis using mdanalysis. Biophysical Journal 2023, 122 (3), 420a–420a.

(52) Naughton, F. B.; Alibay, I.; Barnoud, J.; Barreto-Ojeda, E.; Beckstein, O.; Bouysset, C.; Cohen, O.; Gowers, R. J.; MacDermott-Opeskin, H.; Matta, M.; et al. MDAnalysis 2.0 and beyond: fast and interoperable, community driven simulation analysis. Biophysical Journal 2022, 121 (3), 272a–273a.

(53) Michaud-Agrawal, N.; Denning, E. J.; Woolf, T. B.; Beckstein, O. Software News and Updates MDAnalysis: A Toolkit for the Analysis of Molecular Dynamics Simulations. Journal of Computational Chemistry 2011, 32 (10), 2319–2327. DOI: 10.1002/jcc.21787.

(54) Scherer, M. K.; Trendelkamp-Schroer, B.; Paul, F.; Perez-Hernandez, G.; Hoffmann, M.; Plattner, N.; Wehmeyer, C.; Prinz, J. H.; Noe, F. PyEMMA 2: A Software Package for Estimation, Validation, and Analysis of Markov Models. J Chem Theory Comput 2015, 11 (11), 5525–5542. DOI: 10.1021/acs.jctc.5b00743.

(55) Waskom, M. L. seaborn: statistical data visualization. Journal of Open Source Software 2021, 6 (60), 3021. DOI: 10.21105/joss.03021.

(56) McKinney, W. a. o. Data structures for statistical computing in python; 2010.

(57) Shen, Y.; Bax, A. SPARTA+: a modest improvement in empirical NMR chemical shift prediction by means of an artificial neural network. J Biomol NMR 2010, 48 (1), 13–22. DOI: 10.1007/s10858-010-9433-9.

(58) Kim, D. H.; Wright, A.; Han, K. H. An NMR study on the intrinsically disordered core transactivation domain of human glucocorticoid receptor. BMB Rep 2017, 50 (10), 522–527. DOI: 10.5483/bmbrep.2017.50.10.152.

(59) Kjaergaard, M.; Brander, S.; Poulsen, F. M. Random coil chemical shift for intrinsically disordered proteins: effects of temperature and pH. J Biomol NMR 2011, 49 (2), 139–149. DOI: 10.1007/s10858-011-9472-x.

(60) Kjaergaard, M.; Poulsen, F. M. Sequence correction of random coil chemical shifts: correlation between neighbor correction factors and changes in the Ramachandran distribution. J Biomol NMR 2011, 50 (2), 157–165. DOI: 10.1007/s10858-011-9508-2.

(61) Kabsch, W.; Sander, C. Dictionary of protein secondary structure: pattern recognition of hydrogen-bonded and geometrical features. Biopolymers 1983, 22 (12), 2577–2637. DOI: 10.1002/bip.360221211.

(62) Humphrey, W.; Dalke, A.; Schulten, K. VMD: visual molecular dynamics. J Mol Graph 1996, 14 (1), 33-38, 27-38. DOI: 10.1016/0263-7855(96)00018-5.

(63) Moes, D.; Banijamali, E.; Sheikhhassani, V.; Scalvini, B.; Woodard, J.; Mashaghi, A. ProteinCT: An implementation of the protein circuit topology framework. MethodsX 2022, 9, 101861. DOI: 10.1016/j.mex.2022.101861.

(64) Zhu, J.; Salvatella, X.; Robustelli, P. Small molecules targeting the disordered transactivation domain of the androgen receptor induce the formation of collapsed helical states. Nat Commun 2022, 13 (1), 6390. DOI: 10.1038/s41467-022-34077-z.

(65) Hofmann, H.; Soranno, A.; Borgia, A.; Gast, K.; Nettels, D.; Schuler, B. Polymer scaling laws of unfolded and intrinsically disordered proteins quantified with single-molecule spectroscopy. Proc Natl Acad Sci U S A 2012, 109 (40), 16155–16160. DOI: 10.1073/pnas.1207719109.

(66) Rieloff, E.; Skepo, M. The Effect of Multisite Phosphorylation on the Conformational Properties of Intrinsically Disordered Proteins. Int J Mol Sci 2021, 22 (20). DOI: 10.3390/ijms222011058.

(67) Dahlman-Wright, K.; Baumann, H.; McEwan, I. J.; Almlof, T.; Wright, A. P.; Gustafsson, J. A.; Hard, T. Structural characterization of a minimal functional transactivation domain from the human glucocorticoid receptor. Proc Natl Acad Sci U S A 1995, 92 (5), 1699–1703. DOI: 10.1073/pnas.92.5.1699.

(68) Buholzer, K. J.; McIvor, J.; Zosel, F.; Teppich, C.; Nettels, D.; Mercadante, D.; Schuler, B. Multilayered allosteric modulation of coupled folding and binding by phosphorylation, peptidyl-prolyl cis/trans isomerization, and diversity of interaction partners. J Chem Phys 2022, 157 (23), 235102. DOI: 10.1063/5.0128273.

(69) Hamelberg, D.; Shen, T.; McCammon, J. A. Phosphorylation effects on cis/trans isomerization and the backbone conformation of serine-proline motifs: accelerated molecular dynamics analysis. J Am Chem Soc 2005, 127 (6), 1969–1974. DOI: 10.1021/ja0446707.

(70) Vandevyver, S.; Dejager, L.; Libert, C. Comprehensive overview of the structure and regulation of the glucocorticoid receptor. Endocr Rev 2014, 35 (4), 671–693. DOI: 10.1210/er.2014-1010.

(71) Valsson, O.; Tiwary, P.; Parrinello, M. Enhancing Important Fluctuations: Rare Events and Metadynamics from a Conceptual Viewpoint. Annu Rev Phys Chem 2016, 67, 159–184. DOI: 10.1146/annurev-physchem-040215-112229.

(72) Marsh, J. A.; Forman-Kay, J. D. Sequence determinants of compaction in intrinsically disordered proteins. Biophys J 2010, 98 (10), 2383–2390. DOI: 10.1016/j.bpj.2010.02.006.

(73) Tesei, G.; Trolle, A. I.; Jonsson, N.; Betz, J.; Knudsen, F. E.; Pesce, F.; Johansson, K. E.; Lindorff-Larsen, K. Conformational ensembles of the human intrinsically disordered proteome. Nature 2024, 626 (8000), 897–904. DOI: 10.1038/s41586-023-07004-5.

(74) Iakoucheva, L. M.; Radivojac, P.; Brown, C. J.; O’Connor, T. R.; Sikes, J. G.; Obradovic, Z.; Dunker, A. K. The importance of intrinsic disorder for protein phosphorylation. Nucleic Acids Res 2004, 32 (3), 1037–1049. DOI: 10.1093/nar/gkh253.

