## Supporting Information for "Phosphorylation regulated conformational diversity and topological dynamics of an intrinsically disordered nuclear receptor"

#### Supplementary Information

FASTA sequences of studied phosphovariants:

>NP:

DQSTFDILQDLEFSSGSPGKETNESPWRSDLLIDENCLLSPLAGEDDSFLLEGNSNED

>pSer 203:

DQSTFDILQDLEFSSG[pS]PGKETNESPWRSDLLIDENCLLSPLAGEDDSFLLEGNSNED

>pSer 211:

DQSTFDILQDLEFSSGSPGKETNE[pS]PWRSDLLIDENCLLSPLAGEDDSFLLEGNSNED

>pSer 203-211:
DQSTFDILQDLEFSSG[pS]PGKETNE[pS]PWRSDLLIDENCLLSPLAGEDDSFLLEGNSNED

>pSer 203-211-226:

DQSTFDILQDLEFSSG[pS]PGKETNE[pS]PWRSDLLIDENCLL[pS]PLAGEDDSFLLEGNSNED

### Supplementary Figures

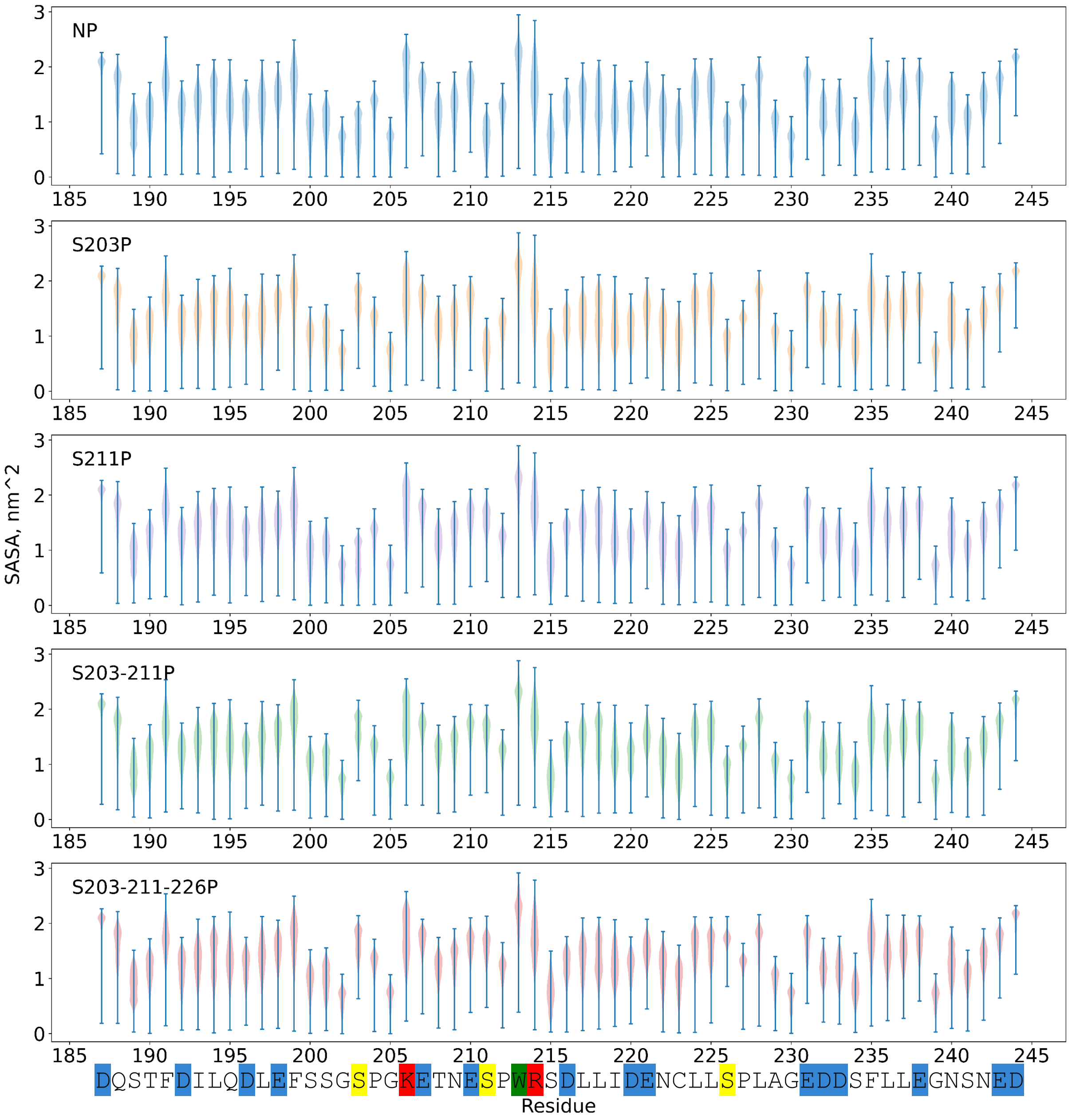


*Figure S2 SASA per redisue. The strong variation and values of SASA higer than 1.5 nm^2 confirm that all studied phoshpovariants were in the disordered state.*

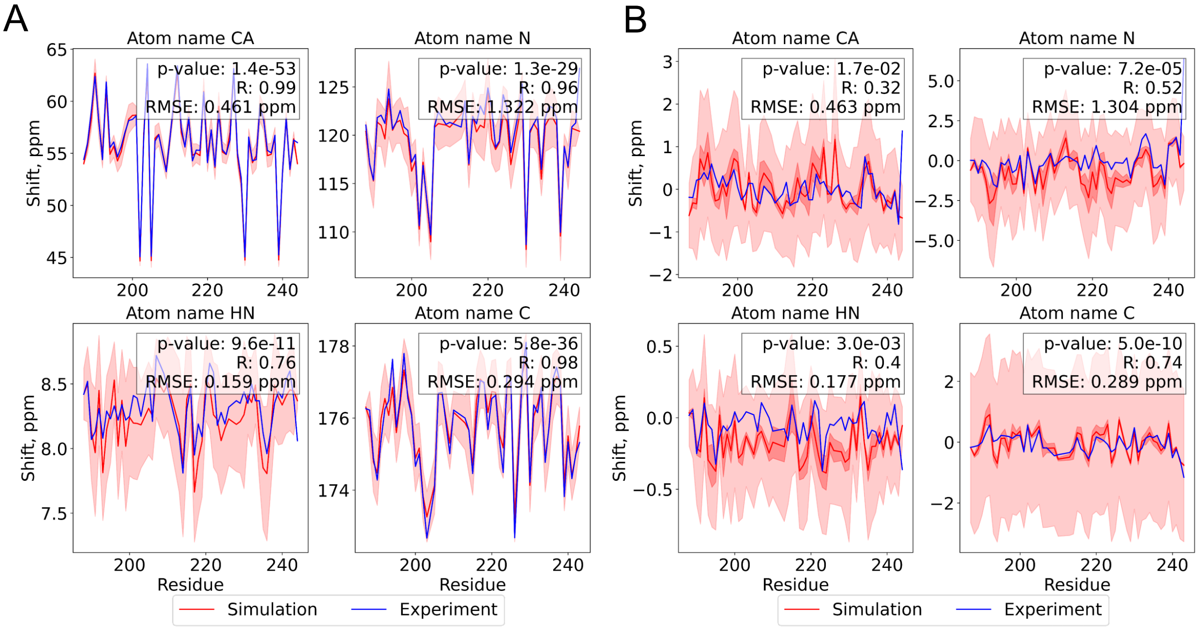


*Figure S3 Computed NMR shifts for non-phosphorylated AF1c (NP) agree well with experimental data [1]. (**A**) Comparison of experimental and calculated NMR shifts. (**B**) Comparison of experimental and calculated NMR shifts corrected to random coil component. For comparison, only backbone atoms shifts were used. Subplots represent the shifts for CA – $^{13}C_{\alpha}$, N – backbone $^{15}N$, HN – $^{1}H$-$^{15}N$ proton, and C – carbonyl chemical shifts. The red line shows the **average** of the NMR signal calculated for an ensemble of structures; the medium red area shows the standard error of the signal; the light red area shows the standard deviation of the signal; the blue line shows the experimentally measured shifts. The Pearson correlation test was calculated between **average** calculated from simulation structures signal (red line) and experimental shift (blue line). p-value is a Pearson test p-value, R is a Pearson product-moment correlation coefficient, RMSE is the root mean squared error. The RMSE was calculated between the average of the calculated signal (red line) and experimental shift (blue line).*

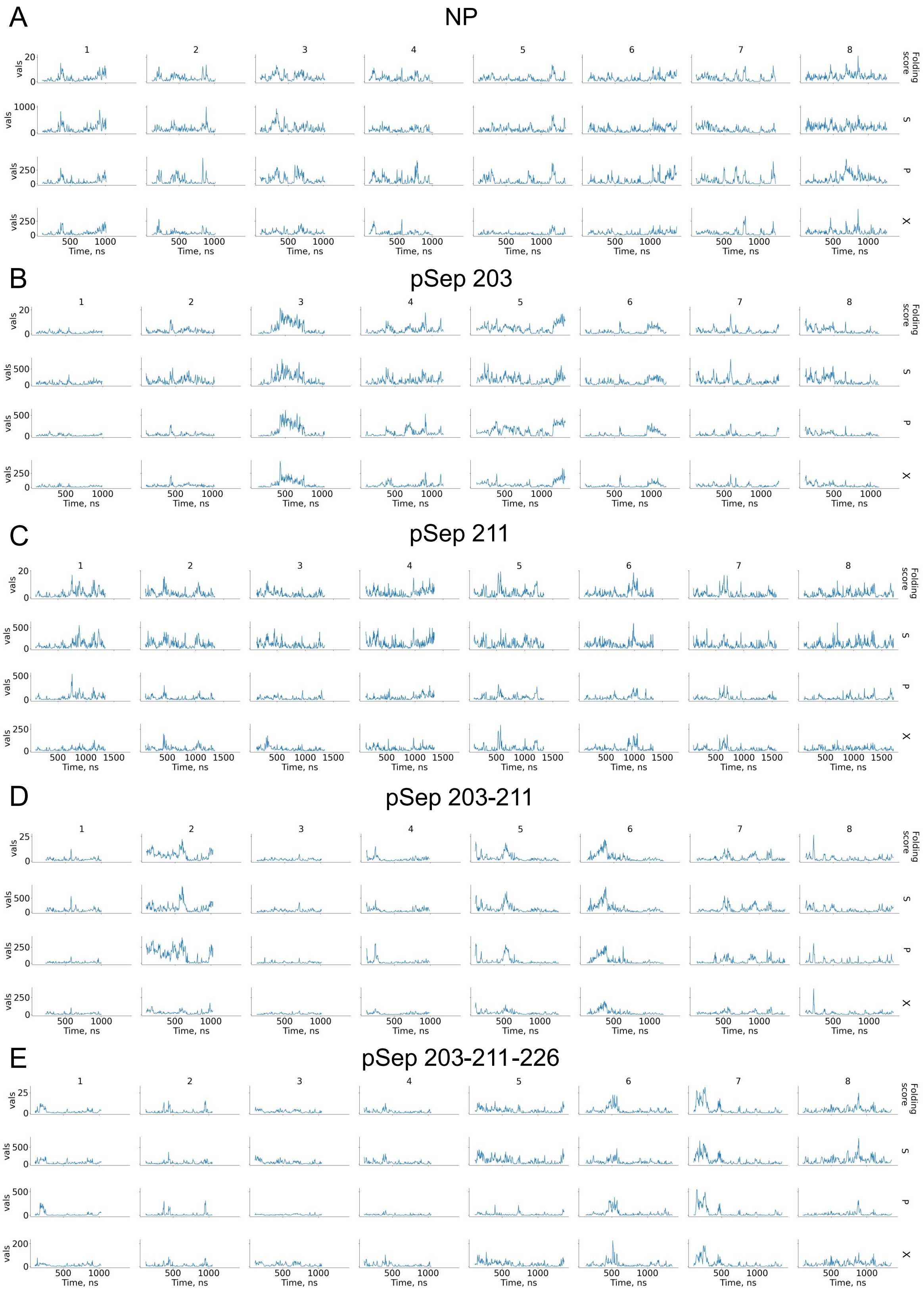


*Figure S4 Protein topology dynamics of different phosphovariants. (**A-E**) Columns represent independent simulations; rows represent various metrics: CT Folding Score, amount of parallel (P), series (S), and cross (X) topological relationships. There is a correlation between all topological relationships, but they are not equal. Significantly, the number of series relationships usually has the biggest amplitude, while cross has the smallest. Thus, for efficient tracking of topological changes, we need a metric that equally combines all relationships, similar to our proposed formulation of CT Folding Score.*

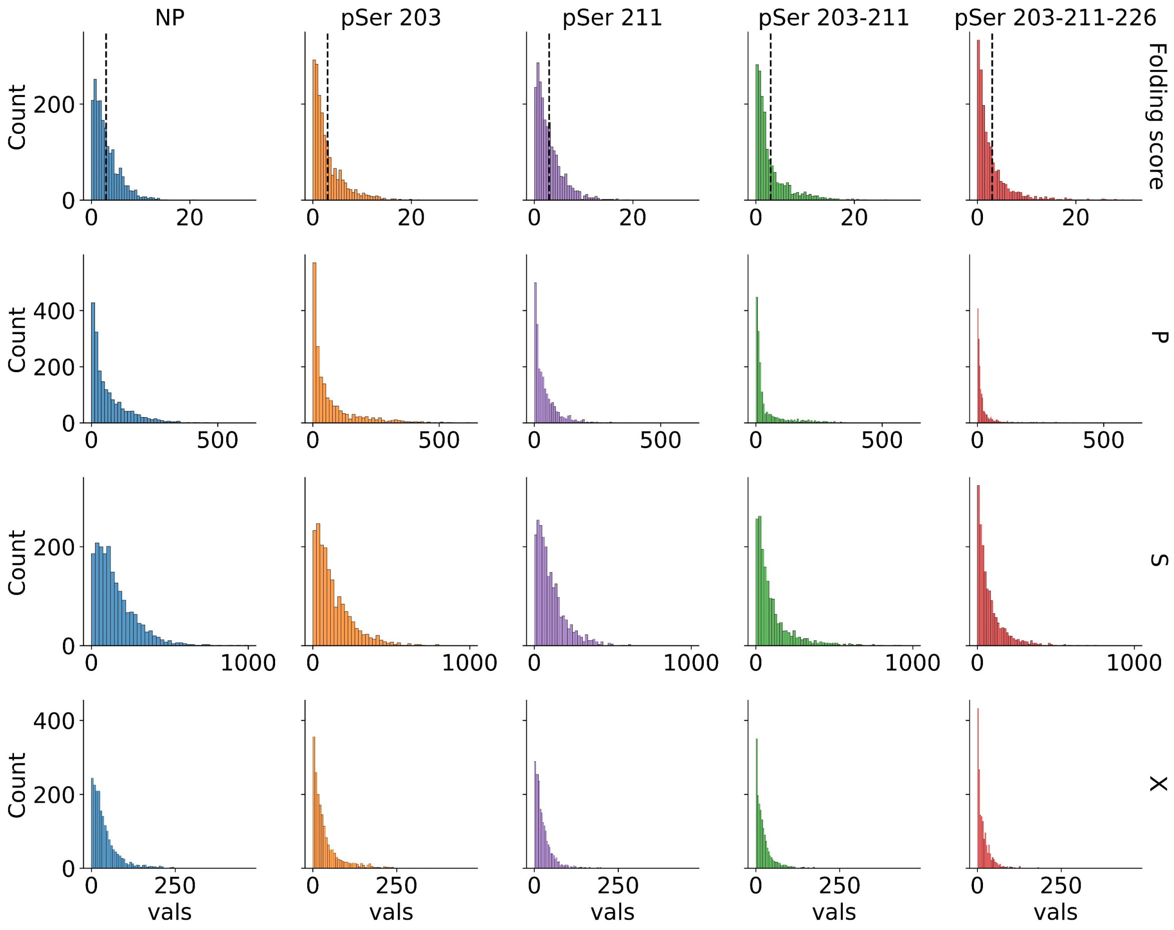


*Figure S5 Histograms of protein topological parameters. Columns denote different phosphovariants, rows represent various metrics: CT Folding Score, amount of parallel (P), series (S), and cross (X) topological relationships. Histograms were constructed after combining the values from independent replicates. Dashed vertical line in histograms for folding score represents threshold. It can be seen that NP phosphovariant has the broadest distribution in P, S, and X values, that correlates with the smallest gyration radius and SASA.*

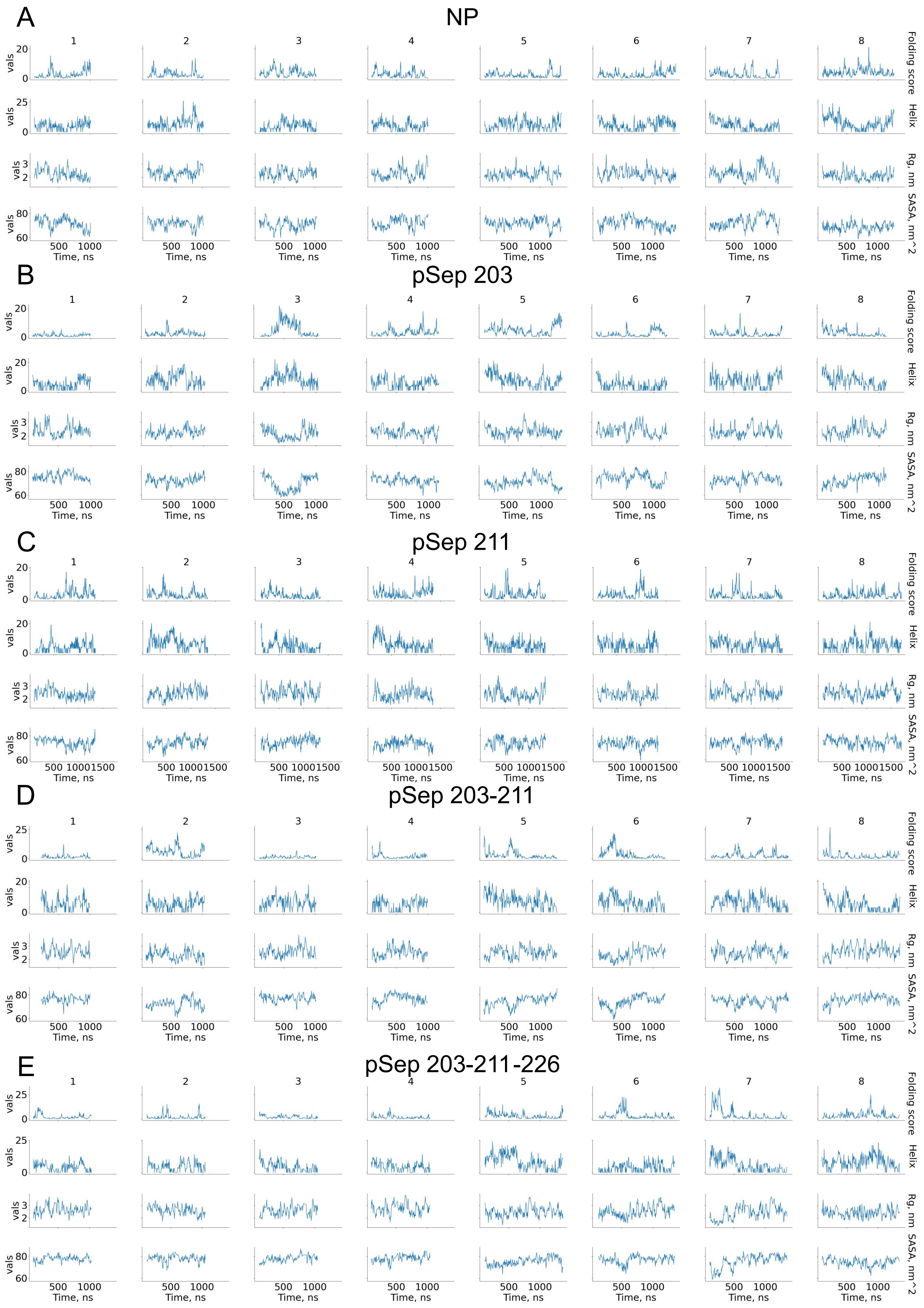


*Figure S6 Dynamics of various protein metrics for different phosphovariants. (**A-E**) Columns represent independent simulations; rows represent various metrics: CT Folding Score, Helix, Rg, and SASA. CT folding Score is highly sensitive to the formation of contacts within the protein structure, giving it a better contrast upon compaction events compared to classical metrics.*

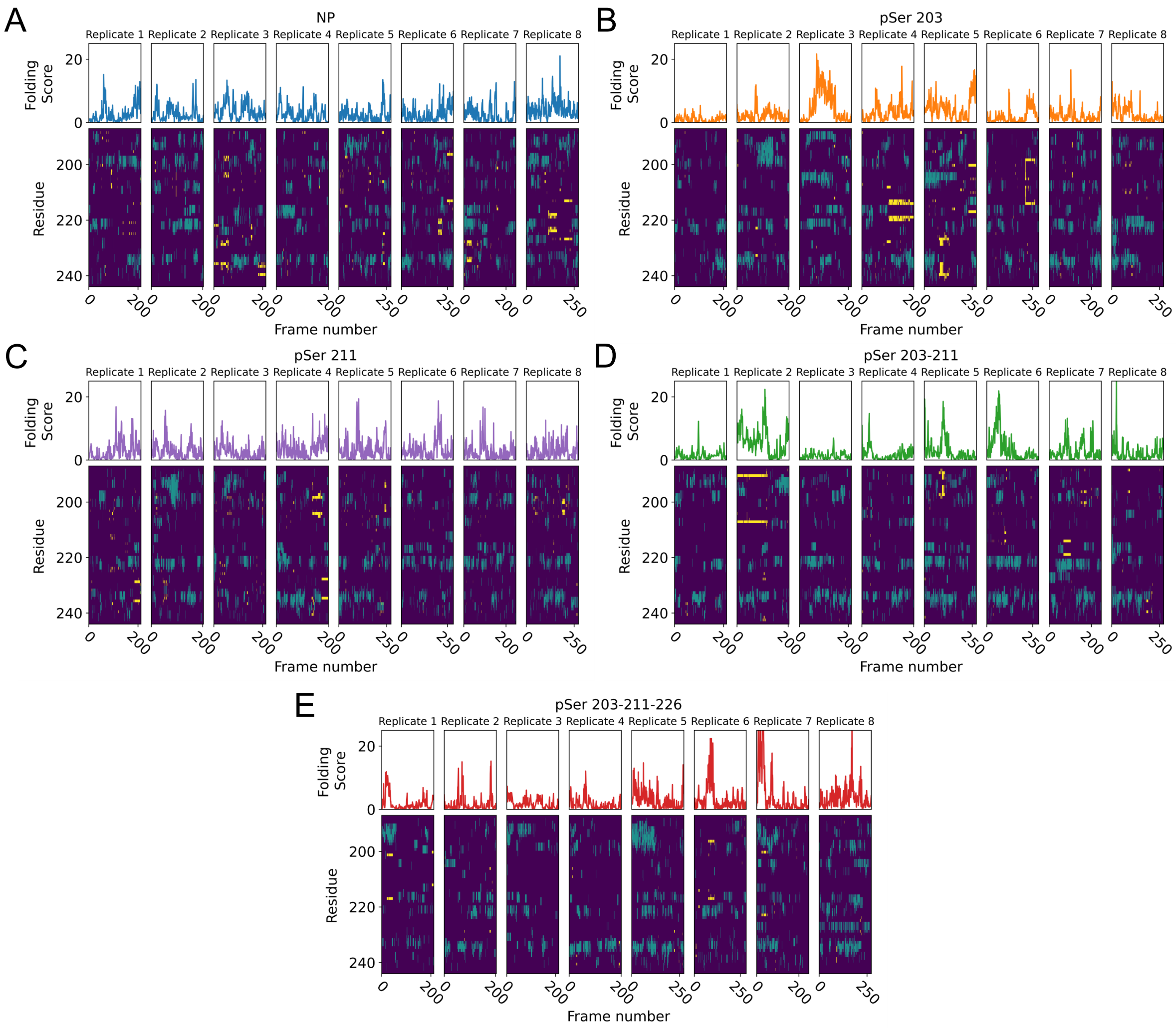


*Figure S7 Dynamics of topology and secondary structure for different phosphovariants of AF1c. (**A-E**) The Folding Score and secondary structure dynamics for different phosphovariants. Top panel – dynamics of the CT Folding Score; bottom panel – dynamics of the secondary structure. Color coding in the secondary structure dynamics plots: dark purple – disordered structure; yellow – elongated structure; green – helical structure.*

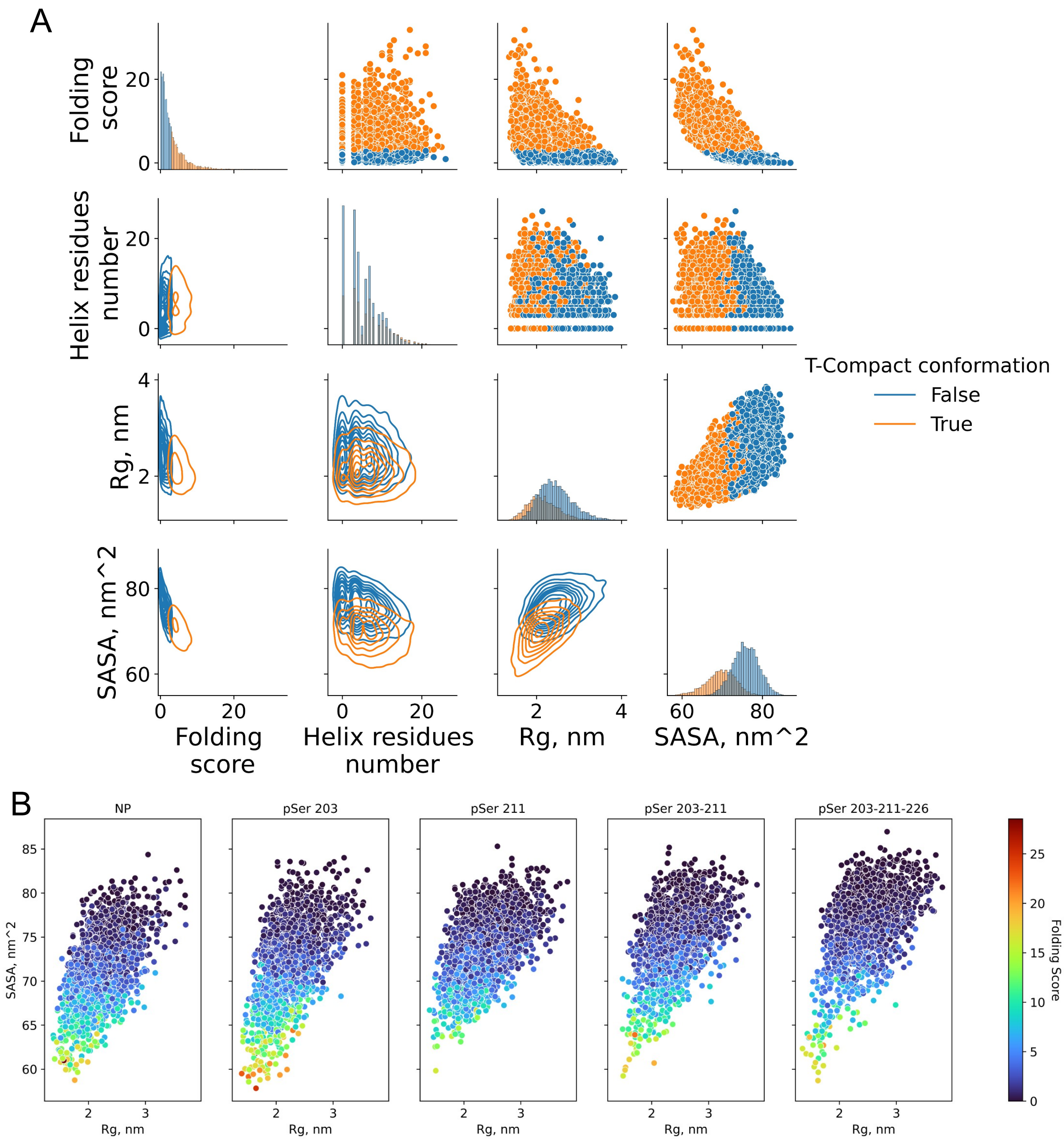


*Figure S8 Relations between CT Folding Score and other protein structure parameters. (**A**) The pairwise relationships between the radius of gyration (Rg), helix residue number, solvent accessible solvent area (SASA), and CT folding score for all phosphovariants combined. On the diagonal are the histograms of CT Folding Score, amount of residues in helix, radius of gyration, and SASA. Colors indicate the distribution of topologically compact conformations (orange) and solvent exposed conformations (blue). In the upper triangle of plots grid, pairwise relationships are shown as scatter plots, while in the lower triangle, the same data is represented using contour plots. (**B**) The distribution of the Rg and SASA for different phosphovariants, colored by the value of the folding score.*

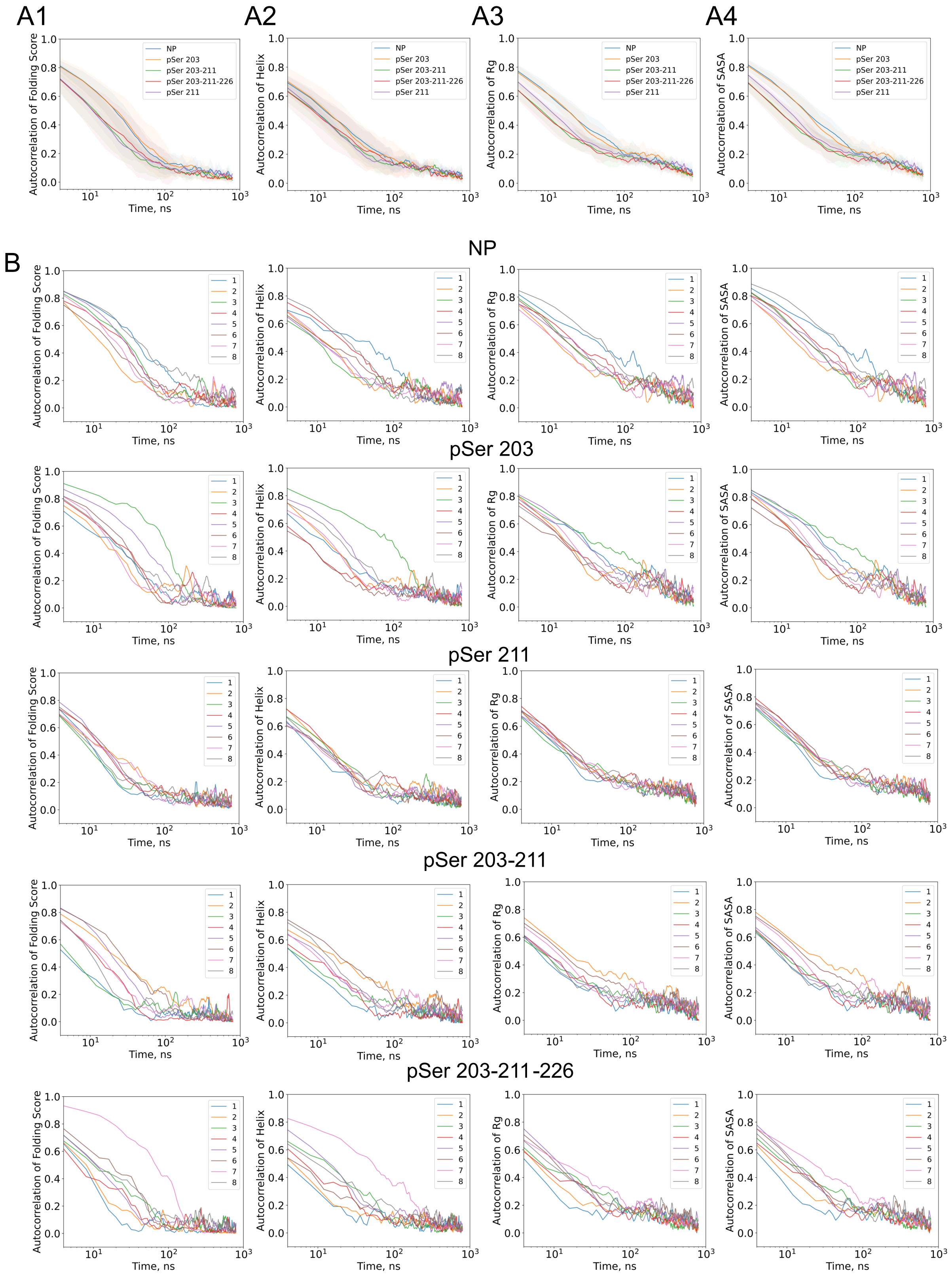


*Figure S9 Autocorrelation of folding score detects differences between different phosphovariant dynamics. Normalized autocorrelation of CT Folding Score (**A1**), number of residues in helical structure (**A2**), Radius of gyration (**A3**), SASA (**A4**). Line represents average between all replicas of autocorrelation, shading represents standard deviation between replicas. (**B**) Autocorrelation of folding metrics for independent trajectories. Columns: CT Folding Score, number of residues in helical structure, radius of gyration, SASA. The data for various phosphorylation variants are grouped by rows.*

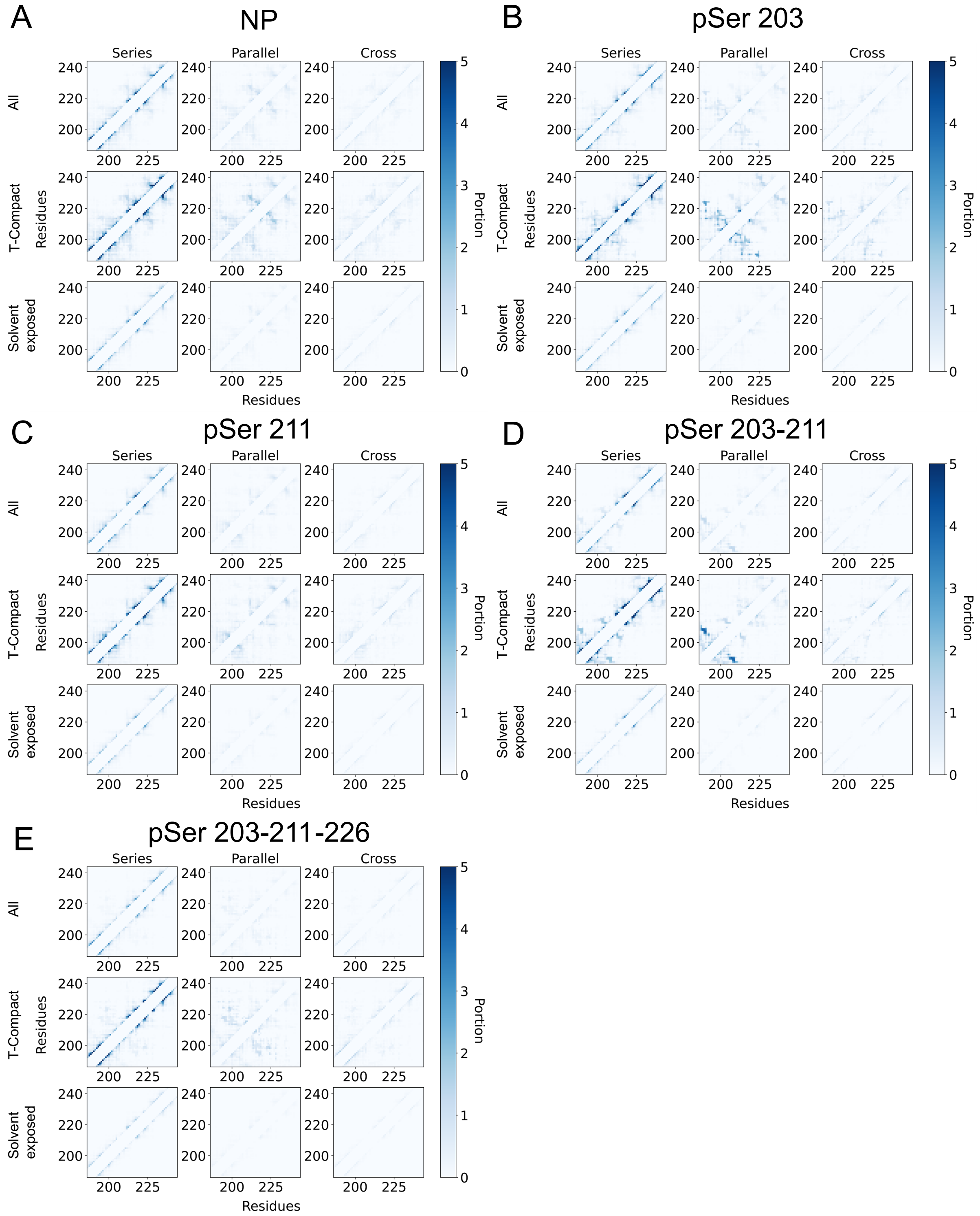


*Figure S10 Analysis of CT matrices of different phosphovariants. (**A-E**) Average amount of topological relations per contact observed during simulations. Color shows the number of topological relations with these contacts participating on average. For this analysis data from all replicates per phosphovariant were combined. In columns, the types of topological relations are shown: Series, Parallel, or Cross. In rows: all, only topologically compacts, or only solvent exposed structures were used for analysis. It can be seen that Series-rich matrices are close to contact matrices, equally considering local and non-local contacts, while parallel-rich matrices are more sensitive to non-local contacts. Cross-rich matrices are sensitive to non-local and contacts in alpha helical structures, as can be seen from the definition of series relationships.*

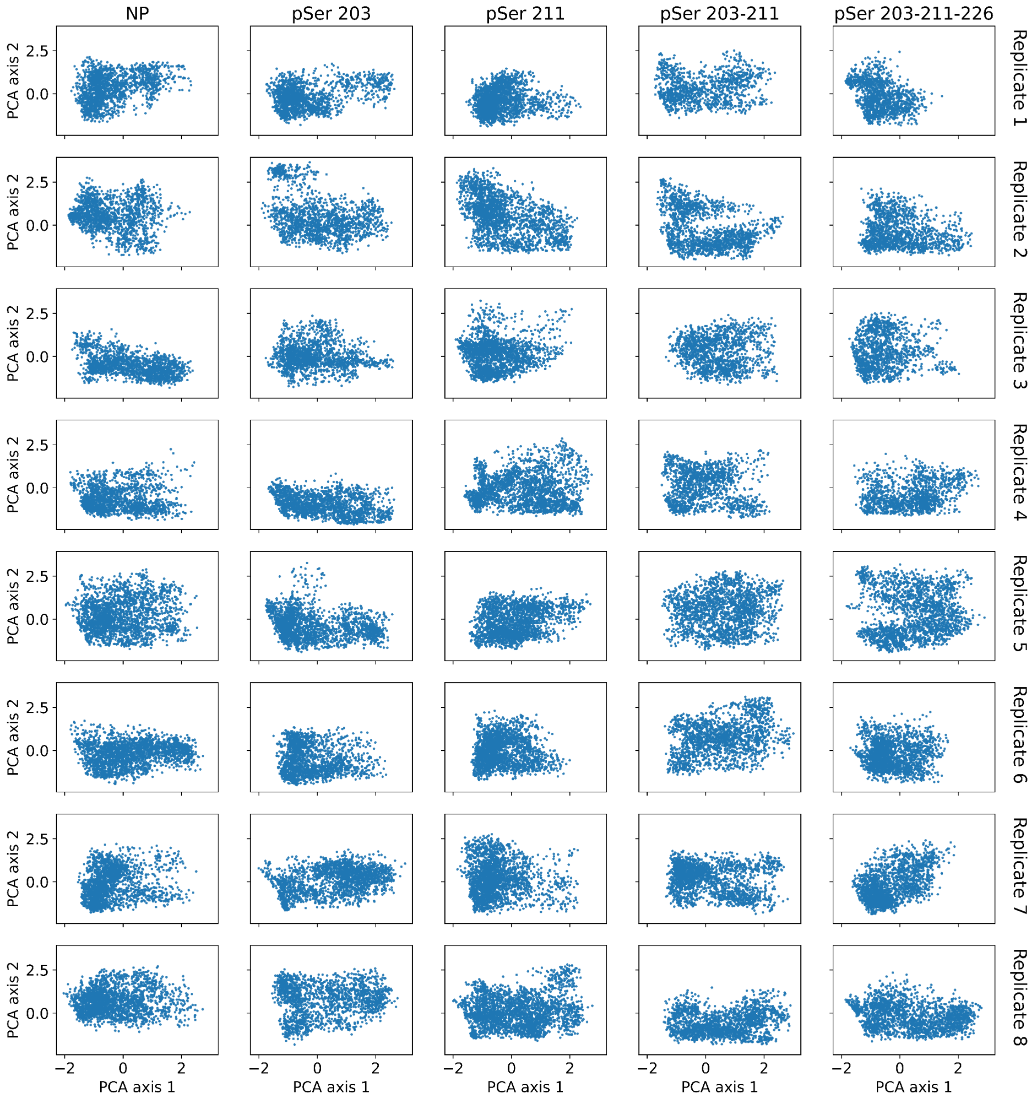


*Figure S11 Dynamics of independent trajectories in PCA axis. Columns represent phosphovariant type; rows represent a number of independent replicates. PCA axis are the same as in Figure 3. It can be seen that none of the simulations were trapped in local conformation, and all trajectories significantly overlap, thus the combined ensemble of structures gives complete statistics for analysis.*

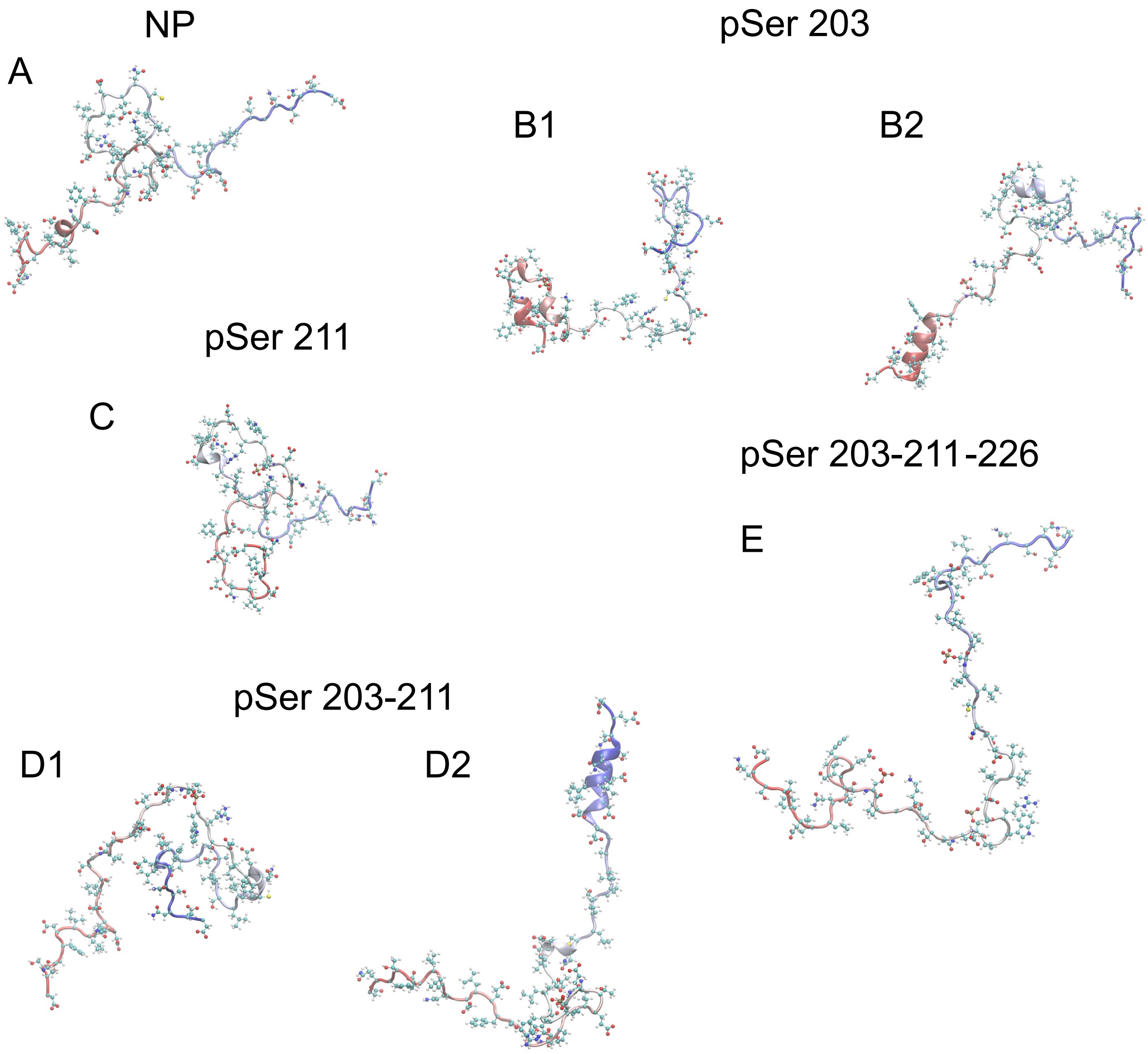


*Figure S12 PCA clustering representative structures. Color indicates the amino acid number: from the red – N-terminal region to the blue – C-terminal region. These structures represent the local free energy minimal shown in Figure 3B. (**A**) Representative structures of the NP molecule. (**B1, B2**) Representative structures of pSer 203 phosphovariant. Panel B1 shows the main conformational state, while B2 highlights a separate minimum featuring an N-terminal helix. (**C**) Representative structure of pSer211. (**D1, D2**) Two representative structures of pSer 203-211, each corresponding to a distinct free energy well. The structure in D2 shows characteristic C-terminal helix. (**E**) The representative structure of pSer 203-11-226. This structure appears solvent-exposed and shows a similarly low level of secondary structure as NP and pSer211. The configurational variability can be seen in Figure 1C.*

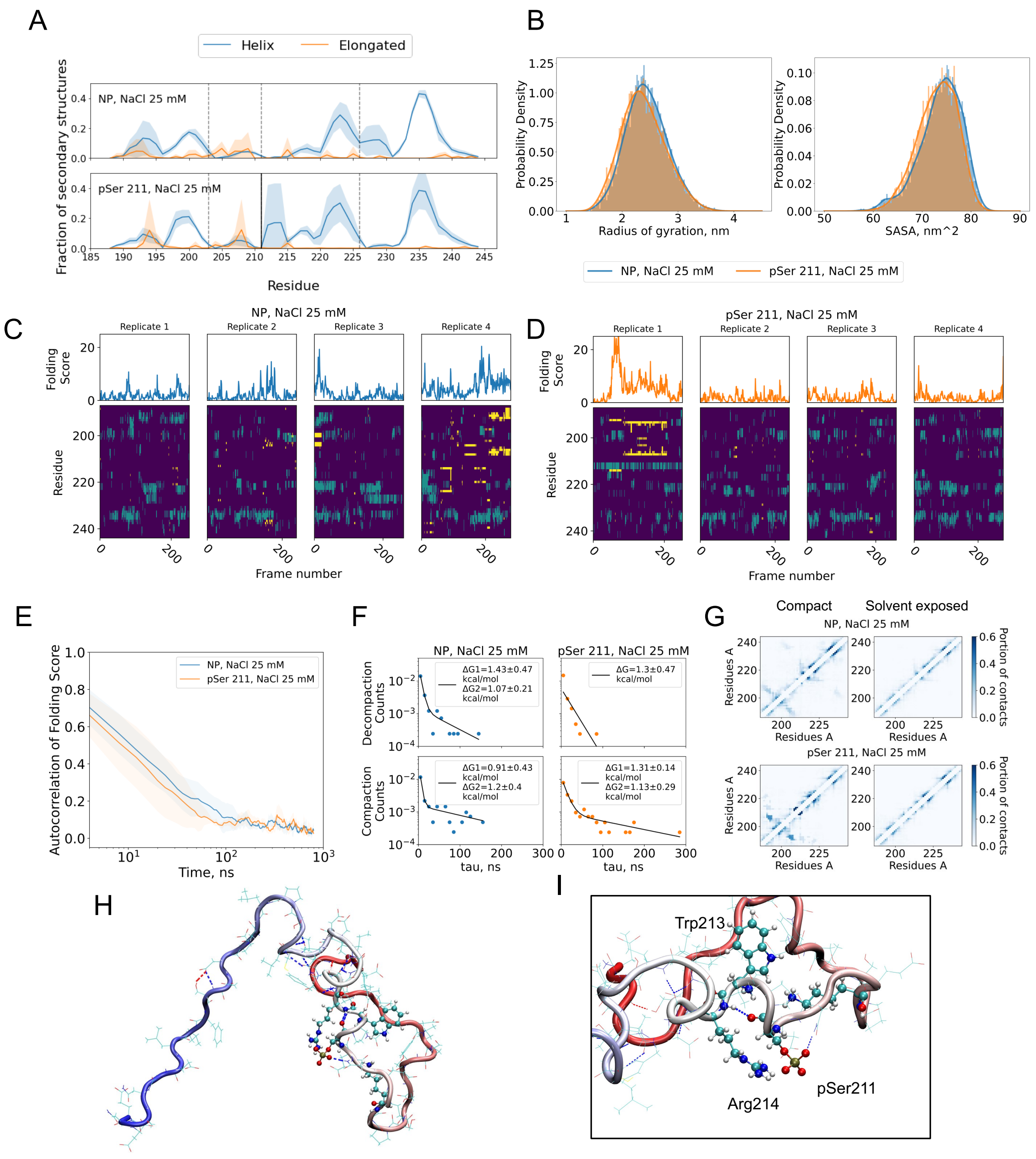


*Figure S13 Results of NP and S211P simulation in low salt condition. In these simulations, protein was simulated with sodium and chloride ions number corresponding to 25 mM NaCl. (**A**) Distribution of secondary structure along the protein chain. (**B**) Distribution of the radius of the gyration and SASA. Interestingly radius of gyration and SASA for both phosphovariants increase in low salt conditions increased, making distributions close to each other. This shows the importance of charge screening for IDP shape. (**C, D**) Dynamics of the CT Folding Score and secondary structure for NP and S211P. Color coding in the secondary structure dynamics plots: dark purple – disordered structure; yellow – elongated structure; green – helical structure. Hight folding score often coexists with entangled structure formation. (**E**) Autocorrelation of CT Folding Score for different phosphorylation variants. NP has some amount of compaction events with 100 ns and 300 ns scale. S211P has compaction events at 60 ns and 300 ns scale. (**F**) Dwell-time histograms fitted with a single exponential decay function. The histograms were constructed from the times protein was in the disordered compact state (Decompaction) and solvent exposed state (Compaction), then counts were divided on the simulation time for the activation energy ΔG calculation. Here, we presented the histogram fits with single and double exponent function; double exponent fits were done on Decompaction times for both NP, and pSer 211 at 25 mM NaCl. (**G**) Contact maps for disordered compact states and solvent exposed states. Interestingly, both phosphovariants show tight*

*contacts in the N-terminal region of the protein. In the contact map of S211P compact state, it can be seen that stable helical region form by pSer211-Arg214 interactions. (**H**) Disordered compact shape of S211P at frame 79 (CT Folding Score is reaching the maximum). Color shows the amino acid number: form red – N-terminal region to blue – C-terminal region. In this image it can be seen that most of the protein is in direct contact with solvent, but there is the contact between N-terminal region and the middle of the protein (compare with D and F). (**I**) Close look on the helix formed by interaction of pSer211-Arg214. This helix is stabilized by charge interaction and backbone hydrogen bond formation (blue bashed line).*

#### References

1. Kim, D.H., A. Wright, and K.H. Han, *An NMR study on the intrinsically disordered core transactivation domain of human glucocorticoid receptor.* BMB Rep, 2017. **50**(10): p. 522-527.
